# Annotating the X-ray diffraction pattern of vertebrate striated muscle

**DOI:** 10.1101/2025.06.11.659175

**Authors:** N. A. Koubassova, D. Dutta, W. Ma, A. K. Tsaturyan, T. Irving, R. Padrón, R. Craig

## Abstract

Low-angle X-ray diffraction is a powerful technique for analyzing the molecular structure of the myofilaments of striated muscle in situ. It has contributed greatly to our understanding of the relaxed, 430-Å-repeating organization of myosin heads in thick filaments in skeletal and cardiac muscle. Using X-ray diffraction, changes in filament structure can be detected on the Å length scale and millisecond time scale, leading to models that are the foundation of our understanding of the structural basis of contraction. As with all X-ray fiber diffraction studies, interpretation requires modeling, which has previously been based on low-resolution knowledge of thick filament structure and is complicated by the contributions of multiple filament components to most X-ray reflections. Here, we use an atomic model of the human cardiac thick filament C-zone, derived from cryo-EM, to compute objectively the contributions of myosin heads, tails, titin, and cMyBP-C to the diffraction pattern, by including/excluding these components in the calculations. Our results support some previous interpretations but contradict others. We confirm that the myosin heads are responsible for most of the intensity on the myosin layer-lines, including the M3 meridional. Contrary to expectation, we find that myosin tails contribute little to the pattern, including the M6 meridional; this reflection arises mainly from heads and other components. The M11 layer line (39 Å spacing) arises mostly from the curved and kinked structure of titin, which allows eleven ∼42-Å-long domains to fit into the 430 Å repeat. The M11 spacing can be used as a measure of strain in the myosin filament backbone as there is negligible head contribution. These insights should aid future understanding of the X-ray pattern of intact muscle in different conditions such as contraction and drug treatment.

**Significance statement:** X-ray diffraction is widely used to study the structure of striated muscle, revealing the molecular organization of the thick and thin filaments in situ. Changes in the X-ray pattern during contraction provide insights into contractile mechanisms on the Å length scale and millisecond timescale. Interpretation of X-ray patterns is based on modeling, which is complicated by contributions of multiple filament components to different reflections and the lack of a reliable thick filament model. Here, we use a cryo-EM-based atomic model of the thick filament to compute contributions of different filament components to the diffraction pattern, by including/excluding these components in the calculations. The insights gained will aid interpretation of the X-ray pattern in relaxation and contraction and following drug treatment.

## INTRODUCTION

Contraction of striated muscle occurs through interactions between overlapping arrays of thick (myosin-containing) and thin (actin-containing) filaments arranged in repeating units called sarcomeres (1–3). Thick and thin filament molecular structures have been elucidated by two complementary techniques: electron microscopy (EM) and X-ray fiber diffraction (3–7). Each method has advantages and disadvantages. EM (negative staining, cryo-EM), combined with three-dimensional (3D) reconstruction, directly reveals the structure of filaments, but these are typically in muscle homogenates, and thus removed from their native environment in the sarcomere (5). X-ray diffraction provides information on filament structure in intact muscle, and can reveal structural changes occurring on the Å length-scale and millisecond time-scale but, like all fiber diffraction studies, depends on modeling for interpretation (6,8,9). Modeling is complicated by the contributions of multiple components to the X-ray reflections and by the lack of a reliable starting model (e.g. (10,11)).

X-ray patterns of relaxed striated muscles consist of actin and myosin “layer lines” coming from the organization of actin and associated components (tropomyosin, troponin) and myosin and its associated proteins (MyBP-C and titin) in their respective filaments (**Fig. 1**, M1-15) (6,12–16). The layer lines have both meridional and off-meridional components. In vertebrate skeletal and cardiac muscle, specific components have been associated with particular reflections. The myosin heads, in crowns of interacting-heads motifs (IHMs), are thought to be the main contributors to the off-meridional 430-Å-based layer lines and to the strong third order meridional reflection (143 Å) (3,6,7,13), reflecting a 430 Å helical repeat and 143 Å axial spacing of IHM crowns. The sixth order meridional (71.5 Å) is thought to be produced by the myosin tails and/or other backbone components, with little or no contribution from the heads in contracting muscle (7,17–19). “Forbidden” meridional reflections occur at orders of 430 Å in addition to those at orders of 143 Å expected for a filament with an axial spacing between myosin crowns of 1/3 the 430 Å helical repeat (3,6,13). These additional reflections are thought to arise from a combination of factors: a deviation from the perfect 143 Å periodicity in the arrangement of myosin heads, and the presence of MyBP-C and titin, which follow a 430 Å axial periodicity (3,6,13,20–22).

**Fig. 1.**
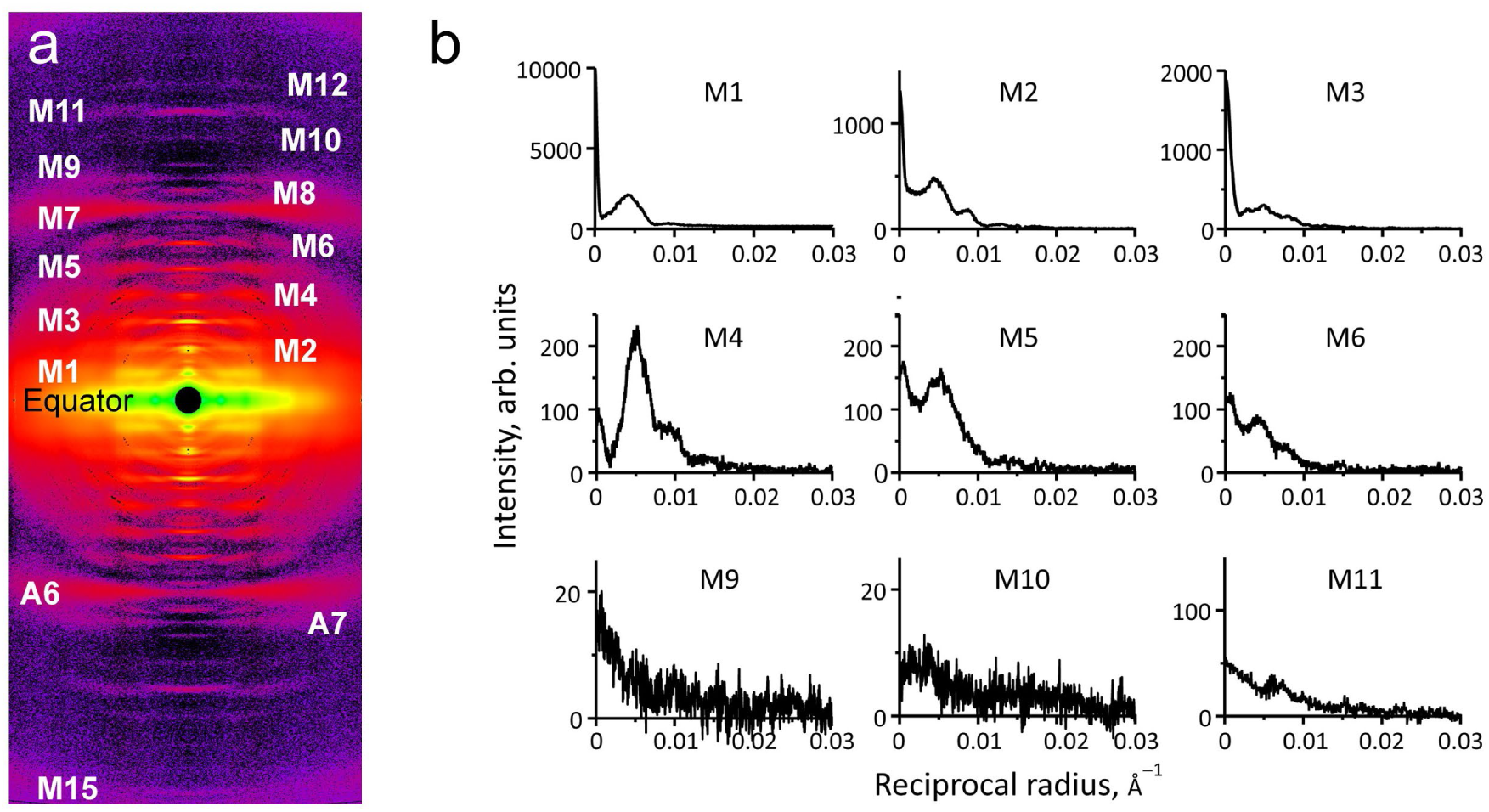
X-ray diffraction of skinned, relaxed porcine ventricular muscle. **(a)** Average of 7 quadrant-folded, background-subtracted X-ray patterns, muscle axis vertical. The pattern exhibits layer lines coming from both actin and myosin filaments that constitute the cardiac muscle sarcomere. Actin layer lines (A6, A7) index on a repeat of ∼355 Å, and myosin layer lines (M1-M15), the focus of this paper, on a repeat of 430 Å. There is no apparent lattice sampling of any of the layer lines, implying that thick filaments have random rotational orientations and/or substantial displacements from their positions in an ideal hexagonal lattice (see text). **(b)** Intensity traces (black) along myosin layer lines after background-subtraction (meridian at left). Layer lines 7, 8 could not be extracted from the pattern due to overlap with thin filament layer lines A6 and A7. The pattern is from muscle treated with mavacamten, for comparison with the pattern computed from the cryo-EM structure, also treated with mavacamten (see text). Note: in patterns recorded on long cameras, some meridional reflections are resolved into multiple finer peaks due to interference between the two halves of the filament (7). The shorter camera used here to capture high angle layer lines, out to M15, does not resolve this detail (see text).

It is now possible to test these assumptions objectively for the first time, using the structure of the relaxed human cardiac thick filament C-zone, which has been solved to 6 Å resolution by cryo-EM (23). Using an atomic model of the C-zone 430 Å repeat based on this structure (PDB 8g4l), we have assessed the likely contributions of the different components to the myosin-based layer lines of the X-ray pattern by computing power spectra – equivalent to the X-ray intensities – of models with different components present or missing. Our calculations demonstrate quantitatively: (1) that the myosin heads are responsible for most of the intensity on the myosin layer-lines, including the M3 meridional (143 Å), as previously suggested; (2) that myosin tails contribute little to the pattern, including the M6 meridional (71.5 Å), contrary to past studies: this reflection arises mainly from myosin heads and possibly other structures outside the C-zone; (3) that titin is the main contributor to the M11 (39 Å) layer line; and (4) that MyBP-C’s contribution depends on whether it lies along the thick filament or extends radially to interact with actin filaments: when extending radially it enhances the forbidden meridional reflections. These insights will aid in understanding the X-ray pattern of intact muscle in future studies, providing a quantitative foundation for interpreting structural changes that underlie models of contraction, and the structural impact of disease and drug treatment.

## MATERIALS AND METHODS

### X-ray diffraction from permeabilized porcine myocardium

Previously frozen heart tissue from male Yucatan pigs was provided by Exemplar Genetics Inc. Humane euthanasia and tissue collection procedures were approved by the Institutional Animal Care and Use Committees at Exemplar Genetics. The procedures were conducted according to the principles in the "Guide for the Care and Use of Laboratory Animals", Institute of Laboratory Animals Resources, Eighth Edition (24), the Animal Welfare Act as amended, and with accepted American Veterinary Medical Association (AVMA) guidelines at the time of the experiments (25). Left ventricular myocardium samples were prepared as described previously (26,27). Briefly, samples were permeabilized in relaxing solution (2.25 mM Na_2_ATP, 3.56 mM MgCl_2_, 7 mM EGTA, 15 mM sodium phosphocreatine, 91.2 mM potassium propionate, 20 mM Imidazole, 0.165 mM CaCl_2_, creatine phosphokinase 15 U/ml, containing 15 mM 2,3-Butanedione 2-monoxime (BDM) and 1% Triton-X100 and 3% dextran at pH 7) at room temperature for 2-3h. Muscles were then washed with fresh relaxing solution for ∼10 minutes, and this was repeated 3 times. The tissue was dissected into ∼300 *μm-*diameter fiber bundles and clipped with aluminum T-clips. The preparations were stored in cold (4°C) relaxing solution with 3% dextran and 50 μM mavacamten for the day’s experiments. X-ray diffraction experiments were performed at the HP BioSAXS & BioSAXS 7A beamline at the Cornell High Energy Synchrotron Source (CHESS). The muscles were incubated in a customized chamber at 28-30°C at a sarcomere length of 2.3 µm in the presence or absence of 50 μM mavacamten (see **Supplemental Discussion: Quantification of the impact of mavacamten on thick filament order**). The X-ray patterns (**Fig. 1**) were collected on an Eiger 4M detector (Dectris, Switzerland) with a 3 s exposure time at HP BioSAXS & BioSAXS.

### X-ray data analysis

X-ray diffraction patterns were converted from *.tiff to *.bsl format and analyzed using BS software (http://muscle.imec.msu.ru/bs_1.htm written by NAK). Seven patterns from three pig hearts without significant tilt and showing high layer line intensities were shifted to a common center, mirrored with respect to the equator and meridian, and then added together to increase the signal-to-noise ratio. Layer line intensities were obtained as described previously (28). Briefly, the intensity was integrated across each myosin layer line in the meridional direction at every reciprocal radius. Background intensity was subtracted using 3-pixel-wide strips on both sides of the layer line, parallel to it. To reduce noise, background intensities were smoothed using a spline function before the subtraction.

### X-ray simulation

Layer line intensities, from M1-M15, were computed from an atomic model (PDB 8g4l) of one 430 Å repeat of the thick filament C-zone, based on a human cardiac thick filament cryo-EM reconstruction (23). The unit cell (**Fig. 2b,c**) included: (1) three 3-fold symmetric crowns of myosin heads (residues 4-836 of the myosin heavy chain (MHC) plus regulatory and essential light chains) in the IHM configuration (29–31), separated axially by ∼143 Å and arranged in a quasi-helix on the filament surface; the 3 crowns had different IHM orientations—Horizontal (CrH), Tilted (CrT) and Disordered (CrD) (23). (2) the filament backbone, containing portions of thirty-three 1590-Å-long α-helical coiled-coil myosin tails (MHC residues > 836) originating in 3-fold symmetric triplets at each of 11 crowns, including the 3 crowns within the repeat and the next 8 more distal crowns. (3) portions of three cardiac myosin-binding protein C (cMyBP-C) molecules and six titin molecules, both arranged along the surface of the backbone. cMyBP-C and titin consist primarily of concatenated immunoglobulin-like (Ig) and fibronectin type 3-like (Fn) domains about 10 kDa in molecular mass and 40-45 Å long (15,16,32,33). Each cMyBP-C in the atomic model comprised 6 domains (domains C5-C10; residues 658 – 1274) of the total 11 cMyBP-C domains in each molecule (34) (domains C0-C4 are disordered and were not part of the model), while each titin molecule consisted of 11 domains (the 11-domain super-repeat of the C-zone that spans the 430-Å repeat (16) (residues 1 – 1084 of C-zone super-repeat 4)). The entire 430-Å repeating structure is built from 3 equivalent, longitudinally oriented sectors (**Fig. 2c**). Nine 430-Å repeats form a complete C-zone in cardiac muscle(35).

**Fig. 2.**
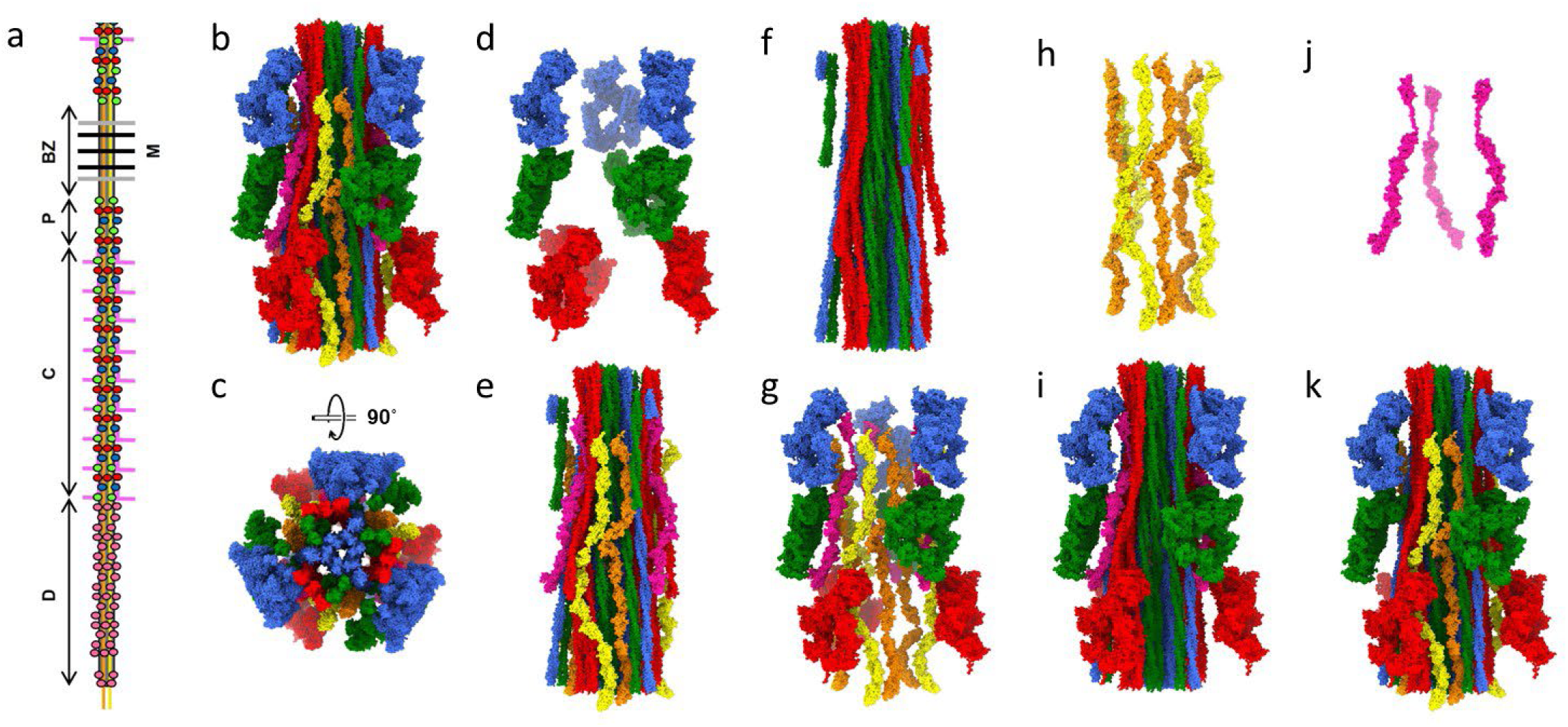
Human cardiac thick filament overall structure and atomic model of 430 Å repeat of C-zone (PDB: 8g4l) showing the individual components alone or removed. **(a)** Cartoon showing P-, C-, and D-zones of half thick filament. Green, red, and blue spheres correspond to crowns CrH, CrD, and CrT, respectively (23). BZ, bare zone; M, M-line. **(b)** 430 Å C-zone repeat with all components, longitudinal view. Crowns with same colors as in **(a)**. Tails originating at the respective crowns have corresponding colors. Titin A and B, orange and yellow respectively, cMyBP-C, pink. **(c)** Full structure in transverse view, viewed towards Z-line. **(d, e)** Same as **(b)** but with heads only **(d)** or heads removed **(e)**. **(f, g)** Same as **(b)** but with tails only **(f)** or tails removed **(g)**. **(h, i)** Same as **(b)** but with titin only **(h)** or titin removed **(i)**. **(j, k)** Same as **(b)** but with cMyBP-C only **(j)** or cMyBP-C removed **(k)**. See also **Fig. S1.**

For optimal computing efficiency, layer line intensities were computed from a single repeat, modeled as being axially repeated an infinite number of times (36). This simplification narrows the layer lines in the meridional direction without affecting the radial distribution of the diffraction intensity. The meridional profile of the diffraction pattern of relaxed muscle was shown to be rich in reflections originating from the C-zone and other parts of the thick filament (13,20,37). Due to the lack of high-resolution structures of the bare zone and P- and D-zones of the myosin filaments (3), we did not model the meridional profiles or the interference between X-rays scattered by the two halves of a filament (19,20,38). These simplifications do not impact our conclusions concerning the major filament contributors to the layer line pattern. The contributions of the different filament components to the X-ray pattern were computed by editing the atomic model in ChimeraX (39) to include or exclude specific components in the calculations. The myosin layer line intensities were calculated as previously described (40) using an all-atom representation, where each protein atom (except hydrogens) had a scattering factor that accounted both for the number of electrons and for surrounding water (41). Mathematical details of the computations are given in Supplemental Information.

The model for radial extension of cMyBP-C domains C0-C6 was constructed by extending these N-terminal domains approximately perpendicular to domains C7-C10, which ran longitudinally as in the atomic model. Residue 870, the N-terminal end of the C7 domain, was taken as the pivot point and a line was drawn through this point perpendicular to the filament axis. Individual cMyBP-C domains were then placed manually, domain by domain, in ChimeraX, so that they were close to that axis and the terminal residues of neighboring domains were in close proximity. Domains C5 and C6 were from PDB 8g4l, C4 and C3 were from AlphaFold prediction AF-Q14896, C2 was from PDB 7lrg, C1 from 7tj7, and C0 from 6cxj.

The model to simulate different degrees of disorder of CrD was produced by assigning different weights (0.0, 0.5, 0.8) to the atoms in the heads of this IHM and computing the resultant layer line intensities. All figures with atomic models were rendered with ChimeraX.

## RESULTS

### Atomic model of the C-zone used in computations

Our goal was to compute the diffraction pattern from the vertebrate thick filament and determine the contributions of its different components (heads, tails, cMyBP-C and titin) to the individual layer lines. Vertebrate filaments comprise seven regions: in each half, there are 3 zones containing myosin heads—a P-zone (*Proximal*), a C-zone (containing *MyBP-C*), and a D-zone (*Distal*)—and at the center a bare zone, containing myosin tails and the M-line, but no heads, where filament polarity reverses (**Fig. 2a**; (3,42)). We used the atomic model of the C-zone 430 Å repeat (PDB 8g4l) as a proxy for the full thick filament in our computations. Our rationale was as follows. The 3,870 Å-long C-zone contains nine 430-Å repeats (35), each consisting of three crowns of quasi-helically ordered myosin heads with ∼143 Å between crowns (a total of 28 crowns per C-zone; (42)), three cMyBP-C molecules and six 11-domain super-repeat titin molecules. The P-zone has only 3 crowns of ordered heads (42) and lacks MyBP-C. The 18 crowns in the ∼2,720 Å-long D-zone (which also lacks cMyBP-C) are thought to be relatively disordered and thus to contribute little to the layer line pattern (14). This is suggested by the absence of any particles resembling the D-zone in our cryo-EM analysis (23), by cryo-ET of thick filaments, which shows a 143 Å repeat (43), but no obvious helical order (P. Luther, personal communication), and by negative staining of thick filaments, which shows only weak ordering of heads in the D-zone compared with the C-zone (RC, unpublished). With only 3 crowns of ordered heads in the P-zone (42), and none in the bare zone, we conclude that the C-zone contains ∼90% [{28/(28+3)} x 100] of the ordered heads in the filament that give rise to the myosin layer lines (and all of the cMyBP-C), and is therefore the dominant contributor to the diffraction pattern and a valid model for its annotation. Using the C-zone 430-Å repeat atomic model (8g4l) as the unit cell, we estimated the contribution of each component (heads, tails, titin, cMyBP-C) to the diffraction pattern by computing the layer line intensities for the full structure and comparing with those from each component alone or from the full structure with the component removed (**Figs. 2d-k, S1c-j)**.

### Myosin heads are the main contributors to the myosin layer lines

We compared the predicted diffraction from the C-zone 430-Å repeat unit containing all components: heads, tails, cMyBP-C and titin (“Full Structure”) with that from the myosin heads alone, and from the full structure with the heads removed (**Figs. 2b, d, e, S1b, c, d)**. For the strongest layer lines (M1-6), the off-meridional intensity from the heads alone was similar to that from the full structure, showing that the heads are the main contributors (**Fig. 3**, magenta *vs.* black lines). This fits with expectations and previous conclusions (6), given that the heads, accounting for ∼45 % of the total mass of the C-zone, form discrete globular structures (IHMs), in contrast with the rest of the mass, consisting of longitudinally extended molecules (tails, cMyBP-C and titin), which would be predicted to contribute primarily to equatorial rather than layer line intensity. When the heads were removed from the full structure, most of the intensity on M1-8 disappeared, consistent with this conclusion (**Fig. 3**, blue *vs.* black). Similar comparisons of the higher order layer lines (M9-15) suggest that components in addition to heads make significant contributions to off-meridional intensity on some layer lines in this weak region of the pattern.

**Fig. 3.**
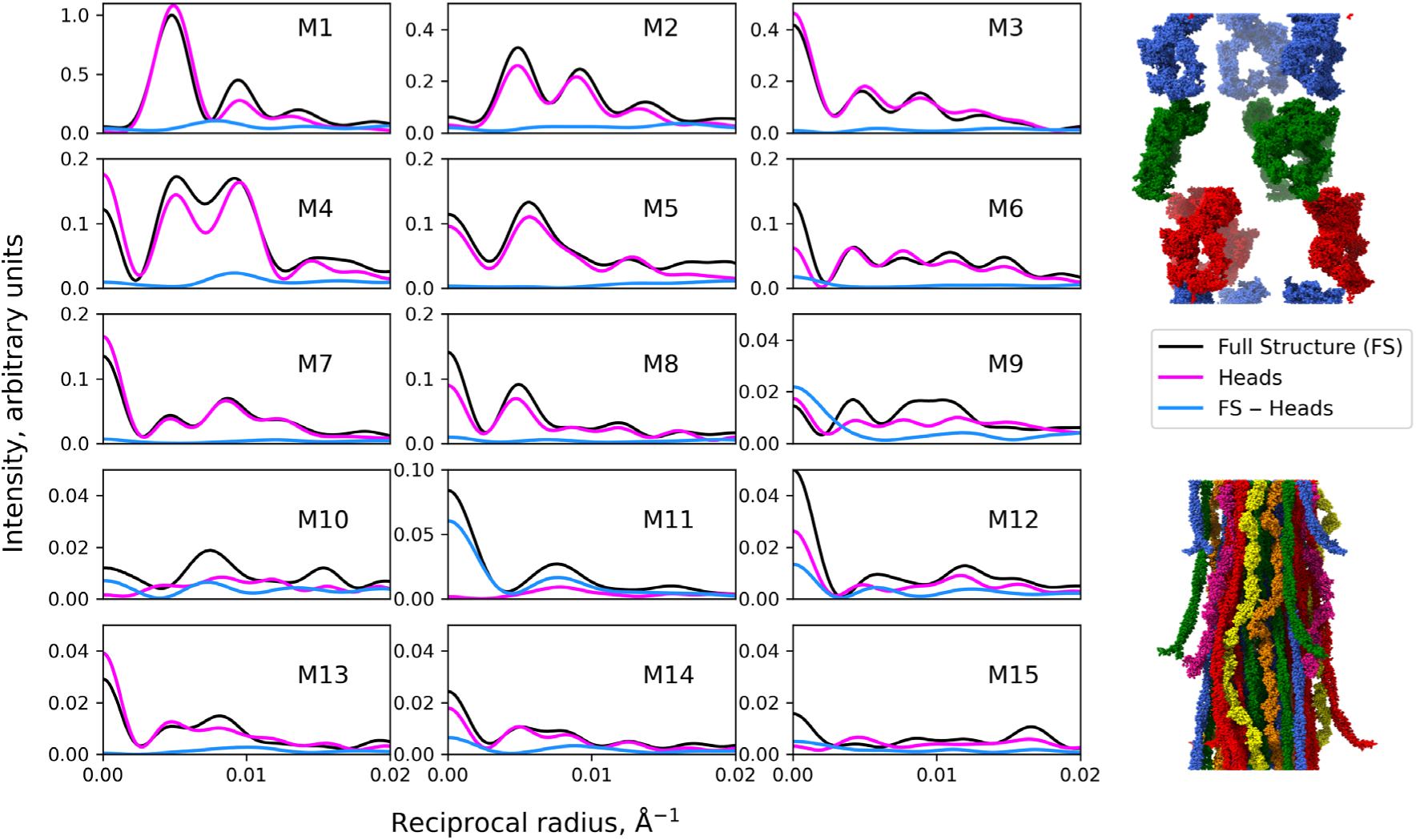
Computed intensities of myosin layer lines 1-15 of cardiac thick filament based on cryo-EM structure PDB 8g4l showing major contribution of heads to the pattern. Black, full structure; magenta, structure with heads only (top right image); blue, structure with heads removed (i.e. backbone only = tails, titin and cMyBP-C; bottom right). The heads account for most of the layer line intensity of the full structure (magenta, blue *vs.* black).

The heads are also seen to be the predominant contributors to the 143 Å meridional reflection (**Fig. 1**; M3): the M3 meridional intensity is strong, similar to the Full Structure, when the heads alone contribute, and comes close to zero when the heads are removed (**Fig. 3**, M3). This is explained by the strong modulation of protein density projected onto the filament axis at intervals of 143 Å coming from the crowns of heads (6,42). The heads also make a major contribution to the M6 meridional reflection (the second order of the M3). With heads alone, the M6 reflection is ∼50% the intensity of the full structure, while when heads are removed the M6 intensity drops to 15% of the Full Structure (**Fig. 3**, M6).

In summary, the pattern from myosin heads alone is similar to the pattern from the full C-zone for the strong layer lines (M1-6), while removal of heads drops most layer lines to near-zero intensity. We conclude that heads are responsible for most of the intensity on these layer lines, including the M3 and M6 meridional reflections.

### Myosin tails contribute little to the myosin layer lines

Myosin tails constitute approximately one half the mass of myosin, but are highly extended parallel to the filament axis. Due to their minimal mass variation/contrast along the filament, they would be expected to contribute little to the myosin layer lines, in contrast with the discrete, globular masses of the myosin heads. We tested the contribution of the tails by comparing layer lines coming from the full structure with those where only tails were present and where tails were removed (**Figs. 2f, g, S1e, f**). Computed diffraction from tails alone showed weak, close to zero, intensity on all layer lines (M1-15), including both meridional and off-meridional components (**Fig. 4**, magenta *vs.* black). Concomitantly, removal of tails (**Fig. 4**, blue *vs.* black) had only a minor impact on the stronger layer lines (M1-6), with a small to moderate effect on M9-10. Strikingly, removal of tails had little effect on the M6 meridional intensity (**Fig. 4**, M6; see **Discussion**). These observations confirm the expectation that myosin tails would make little contribution to the stronger myosin layer lines of relaxed muscle: the myosin heads are the main contributors.

**Fig. 4.**
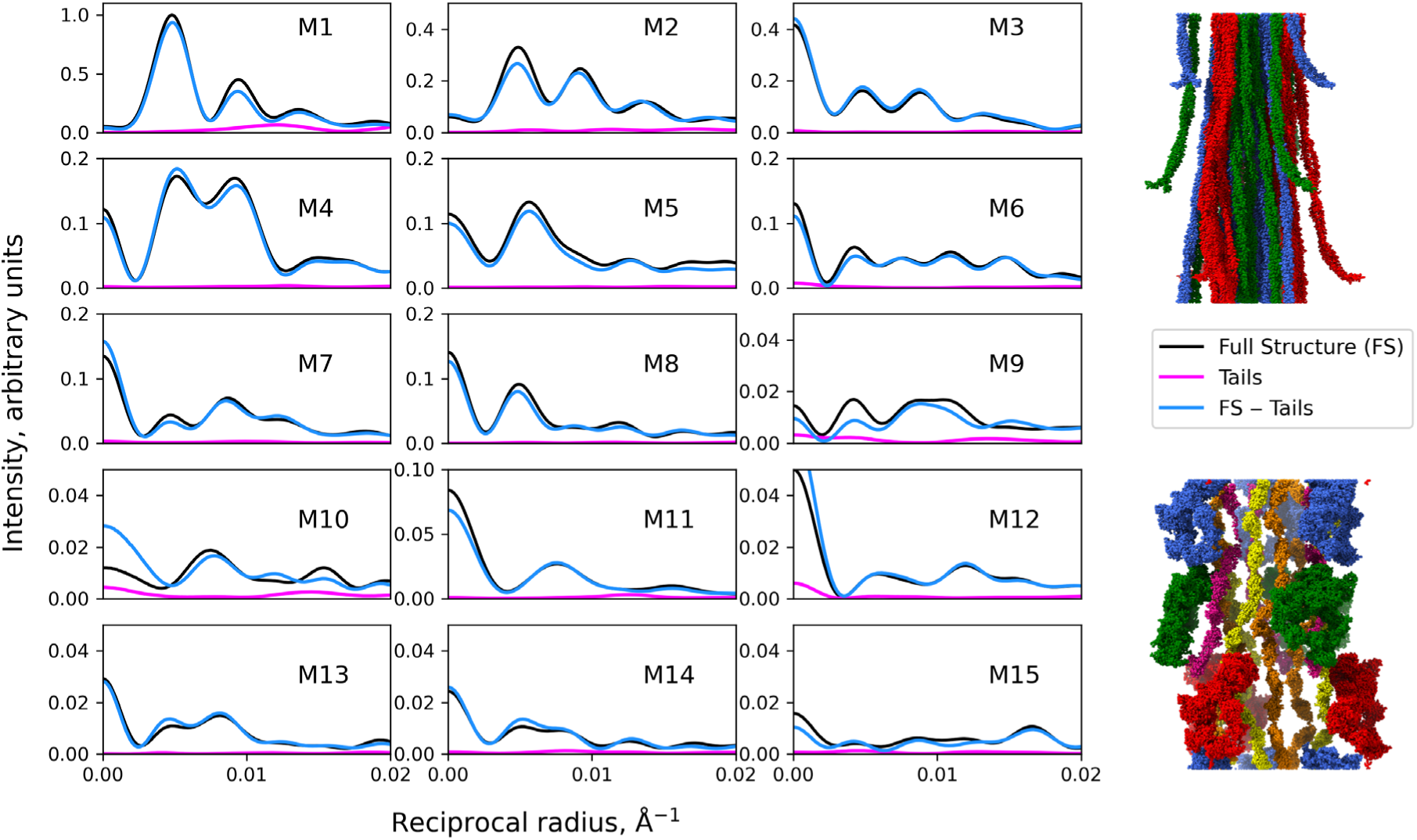
Computed intensities of myosin layer lines 1-15 of cardiac thick filament based on cryo-EM structure PDB 8g4l showing minimal contribution of tails to the pattern. Black, full structure; magenta, structure with tails only (top right); blue, with tails removed (bottom right). The tails contribute little to most layer lines (magenta, blue *vs.* black).

### Titin is the main contributor to the 39 Å layer line (M11)

We investigated the contributions of titin to the diffraction pattern by computing the layer line intensities expected from the 6 titin strands alone with those from the full structure or the full structure lacking titin (**Figs. 2h, I, S1g, h**). Removal of titin had only a small impact on the strong myosin layer lines (M1-6), both meridional and off-meridional components, but a substantial effect on the meridional strength of the weaker layer lines (M10, 11, 12 and 15), where its removal greatly reduced the intensity (**Fig. 5**, blue *vs.* black). Titin alone produced little intensity on any of the strong layer lines (M1-6), but contributed significantly to the weaker part of the pattern (M9-12, magenta *vs.* black; *cf*. (21)). Its strongest contribution was to M11 (the 39 Å layer line), consistent with the near-total (∼11-fold) loss of intensity when titin was removed. This objectively confirms earlier suggestions (21,23,44–46) that titin is the main contributor to the 11^th^ order meridional reflection of the 430 Å repeat pattern, first seen in X-ray diffraction patterns of frog skeletal muscle (6) and optical diffraction patterns of negatively stained A-segments (47). This is explained by the incorporation of eleven 42-Å-long titin domains (the 11-domain super-repeat) into the 430 Å repeat of the filament, made possible by the curving and kinking of the six titin strands as they pass along the filament surface (23).

**Fig. 5.**
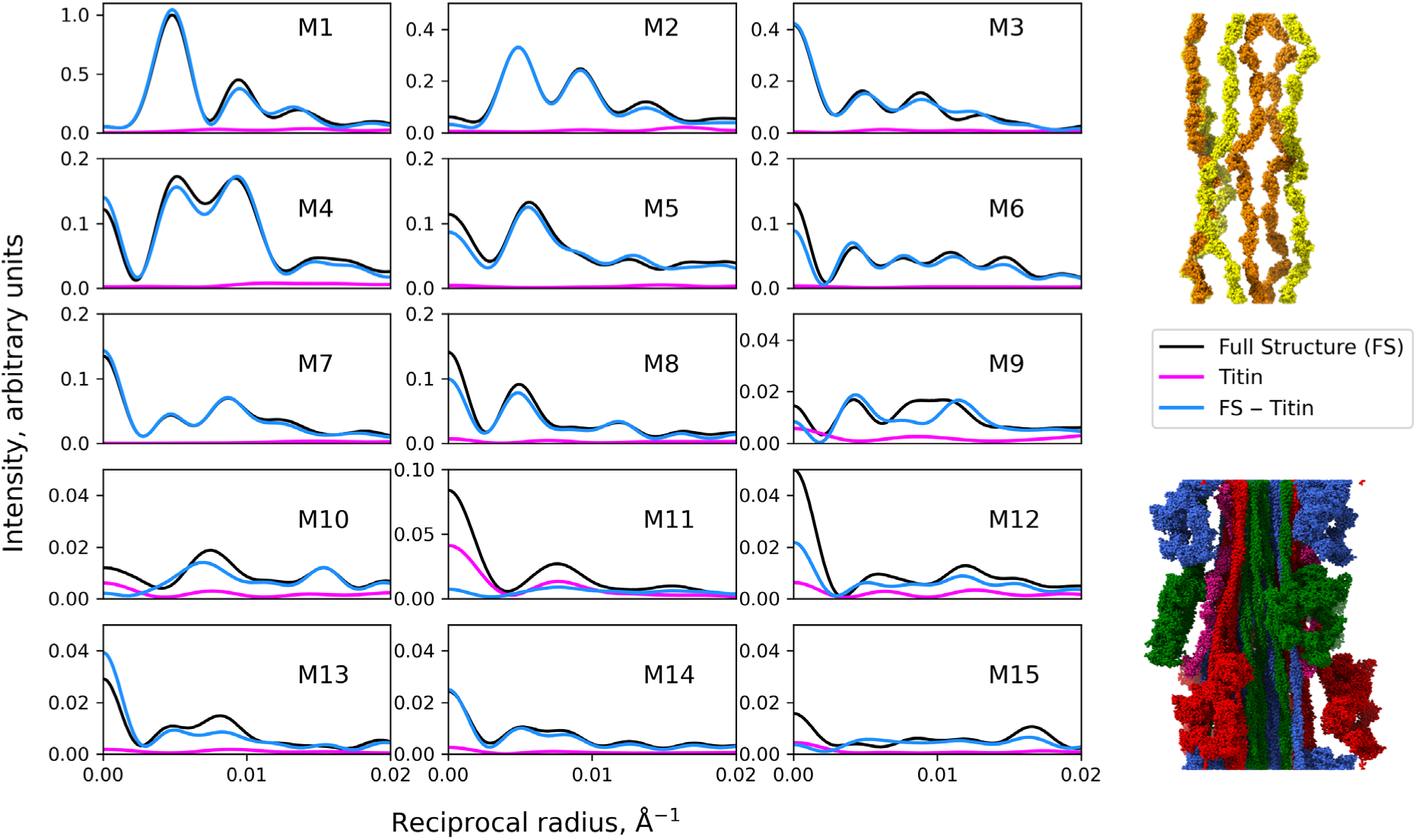
Computed intensities of myosin layer lines 1-15 of cardiac thick filament based on cryo-EM structure PDB 8g4l showing major contribution of titin to M11. Black, full structure; magenta, structure with titin alone (top right); blue, with titin removed (bottom right). Titin contributes primarily to M11 (magenta, blue *vs.* black).

### MyBP-C’s C-terminal half contributes little to the diffraction pattern

The C-terminal half of cMyBP-C in the atomic model (domains C5-10, pink in **Fig. 2b**) extends axially along the filament surface, from contact of the C10 domain with the free head of crown CrT, through C8 and C5 interaction with the free head of the next crown distal to the M-line (CrH) (23). The cryo-EM map also showed domains C2-C4 (towards the N-terminal end) associating with the next most distal crown (CrD), but these were weaker and only visible at low contour, suggesting that they are relatively mobile (23). The C2-C4 domains were therefore not included in the atomic model. The N-terminal region (C1-M-C0) did not appear as discrete domains, suggesting high mobility, and was also omitted from the atomic model.

We investigated the contributions of cMyBP-C domains C5-C10 to the diffraction pattern by comparing the layer line intensities expected from the full atomic model with those from these domains alone, and from the full model lacking them (**Figs. 2j, k, S1i, j**). C5-C10 alone contributed almost no intensity to either the strong or the weak layer lines (both meridional and off-meridional), except for a small intensity on the weak M10 meridional reflection (**Fig. 6**, magenta *vs.* black). Consistent with this, removal of these domains had minimal impact, except for a substantial fractional effect on M10 (**Fig. 6**, blue *vs.* black). The contribution of cMyBP-C specifically to M10 (43 Å reflection) is consistent with the ∼42 Å spacing of Fn and Ig domains in the straight, extended organization of these domains seen in cMyBP-C (23), in contrast with the kinked and curved organization of similar Fn and Ig domains in titin, which reduces their average spacing to ∼39 Å, as discussed in the previous section.

**Fig. 6.**
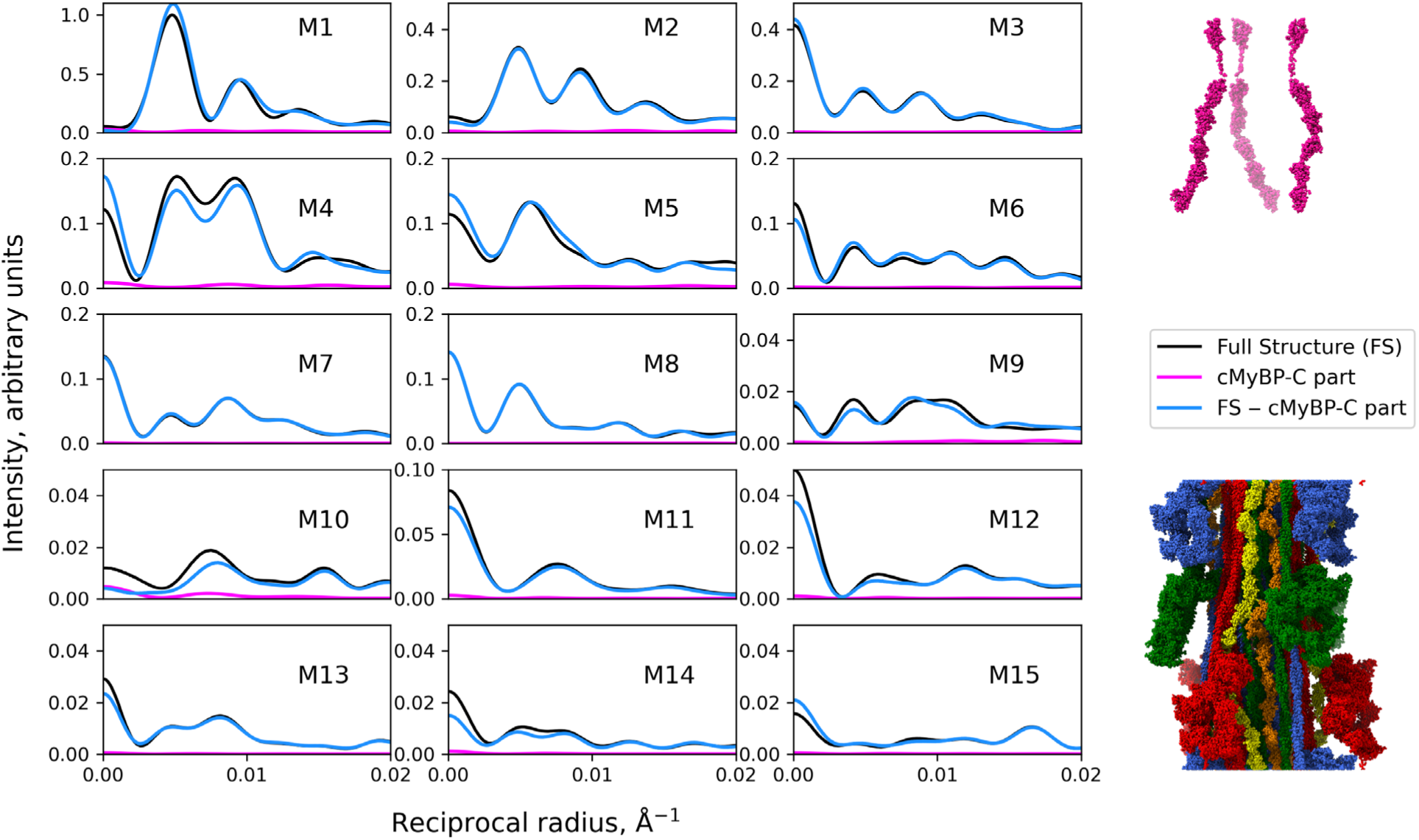
Computed intensities of myosin layer lines 1-15 of cardiac thick filament based on cryo-EM structure PDB 8g4l showing minimal contribution of cMyBP-C C-terminal domains C5-C10 to layer lines. Black, full structure; magenta, structure with cMyBP-C C5-C10 alone (top right); blue, with cMyBP-C C5-C10 removed (bottom right).

The weakness of cMyBP-C’s contributions to the predicted diffraction pattern parallel those of titin and the myosin tails, also with extended structures. Thus the C-terminal half of cMyBP**-**C, with its low mass and its longitudinally extended structure in isolated filaments, would contribute little to the thick filament diffraction pattern. Below, we consider the effect on the pattern of extension of cMyBP-C’s N-terminal region from the thick filament to actin filaments that occurs in the intact sarcomere (42,48).

### Origin of forbidden meridional reflections

For a filament with a strictly helical, 430-Å repeat of myosin molecules, with crowns axially spaced by one third of this (143 Å), and no other components, intensity will occur on the meridian only at every third order of 430 Å (M3, 6, 9, etc.). This is because the 143 Å repeat projected onto the filament axis, which effectively generates the meridional pattern, is exactly 1/3 the helical repeat. While these reflections are clearly observed in experimental X-ray patterns of vertebrate skeletal and cardiac muscle in the relaxed state, meridional intensity is also seen on other myosin layer lines (M1, M2, M4, M5, etc.) (6,7,49) (**Fig. 1**). This suggests regular axial perturbations from the 143 Å periodicity of myosin heads or tails (quasi-helicity) and/or the presence of other components, such as titin and cMyBP-C with a 430-Å-repeat. As a result, projected protein density would exhibit a repeat of 430 Å, giving rise to additional, “forbidden” meridional reflections (3,6,13,21). Although the full meridional profile requires more detailed and complex modeling, our approach allows us to estimate the approximate contribution of these factors.

### Contribution of myosin heads to forbidden meridionals

The IHMs in the cryo-EM map, and in the resulting atomic model, have different tilts in the different crowns of the 430 Å repeat (CrH, CrT and CrD; **Fig. 2b**), and are also displaced from the exact 143 Å axial spacing and 40° azimuthal rotation between crowns required for a perfect helix (23). The intensities calculated from the heads-only structure show that these perturbations from helicity could contribute substantial forbidden meridional intensity on M4, M5, M7, M8, M13 and M14 (**Fig. 3**, magenta). However, the strongest forbidden meridionals (M1 and M2; **Fig. 1**) are not explained in this way.

In addition to the differing tilts and positions of IHMs in the 430 Å repeat, crown CrD is weaker and noisier than CrH and CrT, suggesting some mobility (dynamic disordering) of these heads (23). Different degrees of mobility can be simulated to a first approximation by giving different weights to CrD from 0.0 (full disorder) to 1.0 (fully ordered) (**Fig. 7**, see **Materials and Methods**). This weakened the off-meridional intensity of the strong layer lines (M1-6) compared with the full structure, as expected from the reduction of diffracting head mass (**Fig. 7**, magenta, blue dashed, orange dashed *vs.* black). In contrast, there was an *increase* in intensity of the *meridional* reflections M1, M2, and M5 when CrD heads were removed (weight 0.0) or weighted less (**Fig. 7**, meridional regions). Thus, full or partial disordering of CrD heads could in principle contribute substantially to creation of the strong forbidden meridionals at M1 and M2 although it would have little impact on higher order forbidden meridionals (M11), or even weaken them (M4). The intensity increase of the low order meridionals (M1, M2) is readily explained because reduced weighting or removal (partial or full disordering) of every third crown strongly enhances the perturbation (at low resolution): the projected mass onto the filament axis now has a strong 430 Å density fluctuation compared with the more subtle perturbations when CrD is present at full weight. Although the extreme of full disordering of heads leads to a rise of M1 and M2 meridional intensity similar to that observed experimentally, we believe disordering is only partial in intact muscle (see **Discussion: Explaining the forbidden meridional reflections**).

**Fig. 7.**
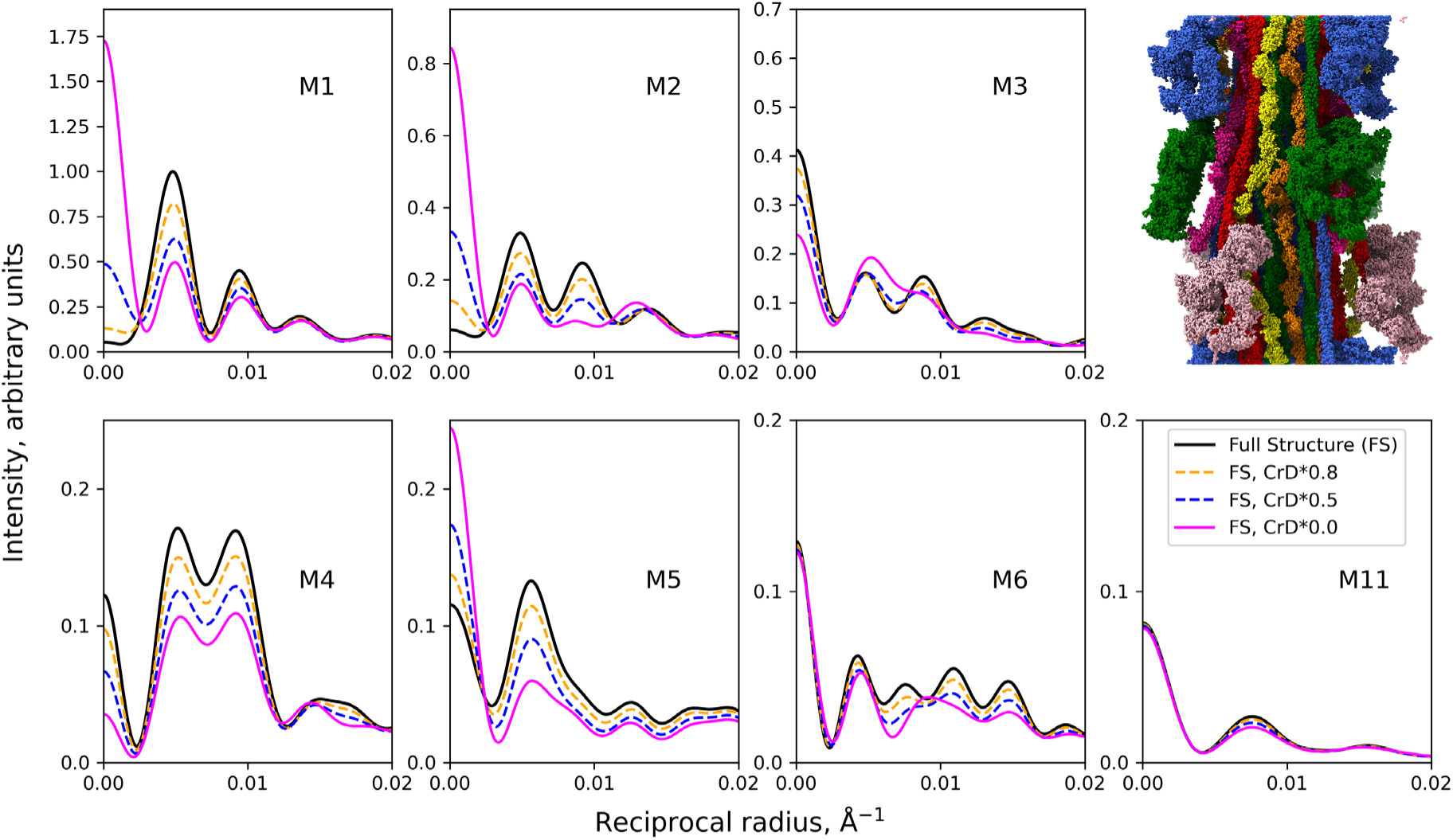
Computed intensities of myosin layer lines 1-6 and 11 from cardiac thick filament based on cryo-EM structure PDB 8g4l showing effect of disordering CrD to different degrees. Black, full structure (CrD fully ordered; top right image); orange, blue, magenta, CrD heads at 0.8, 0.5 or 0.0 weights, simulating CrD disorder of 20%, 50% or 100%, respectively. Full disorder of CrD heads leads to strong contributions to forbidden meridional reflections M1, M2, and M5 (magenta curves), while intermediate levels of disorder have varying impacts, generally lowering the off-meridional intensity in proportion to the degree of disorder.

### Contribution of cMyBP-C to forbidden meridionals

Our atomic model is based on cryo-EM of isolated thick filaments, i.e., filaments removed from the native lattice of the myofibril (23). It thus shows thick filament molecular organization in the absence of potential interactions with actin filaments or other sarcomeric components. In the isolated filament atomic model, domains C5-10 of cMyBP-C, longitudinally organized as described above, have little impact on the filament power spectrum, including the meridian (**Fig. 6**). However, in the intact sarcomere, the more N-terminal domains (roughly C0-C6), mostly absent from the atomic model due to disorder, project from the thick filament at 430 Å intervals, either perpendicular to the filament axis (48), or at varied angles centered on the perpendicular (42), and bind to the thin filament.

The strongly oscillating, 430-Å repeating density created by this transverse organization of domains C0-C6 of cMyBP-C would be expected to contribute significantly more to meridional reflections at orders of 430 Å, at both allowed and forbidden positions, than in our computations based only on the longitudinally arranged C-terminal domains in the isolated filament. We can estimate the magnitude of cMyBP-C’s potential contribution to the forbidden meridional reflections with C0-C6 in this transverse orientation by moving domains C5 and C6 and adding domains C0-C4 so that the N-terminal half of the molecule (C0-C6) projects perpendicular to the filament and comparing the resulting meridional intensities with those from the full structure already described, where domains C5-C10 are longitudinal (**Fig. 8**) and C0-C4 are missing. The results show that when the N-terminal half of cMyBP-C is perpendicular, the M1 and M2 meridionals are both substantially enhanced (**Fig. 8**, magenta and blue *vs.* black). On the other, weaker, forbidden meridionals, there is little impact (M4) or weakening of the intensity (M5, **Fig. 8**, magenta and blue *vs.* black). Thus, the meridional intensities are highly sensitive to the orientation of the C0-C6 or C3-C6 domains, with the larger number of domains (C0-C6) having the greater impact.

**Fig. 8.**
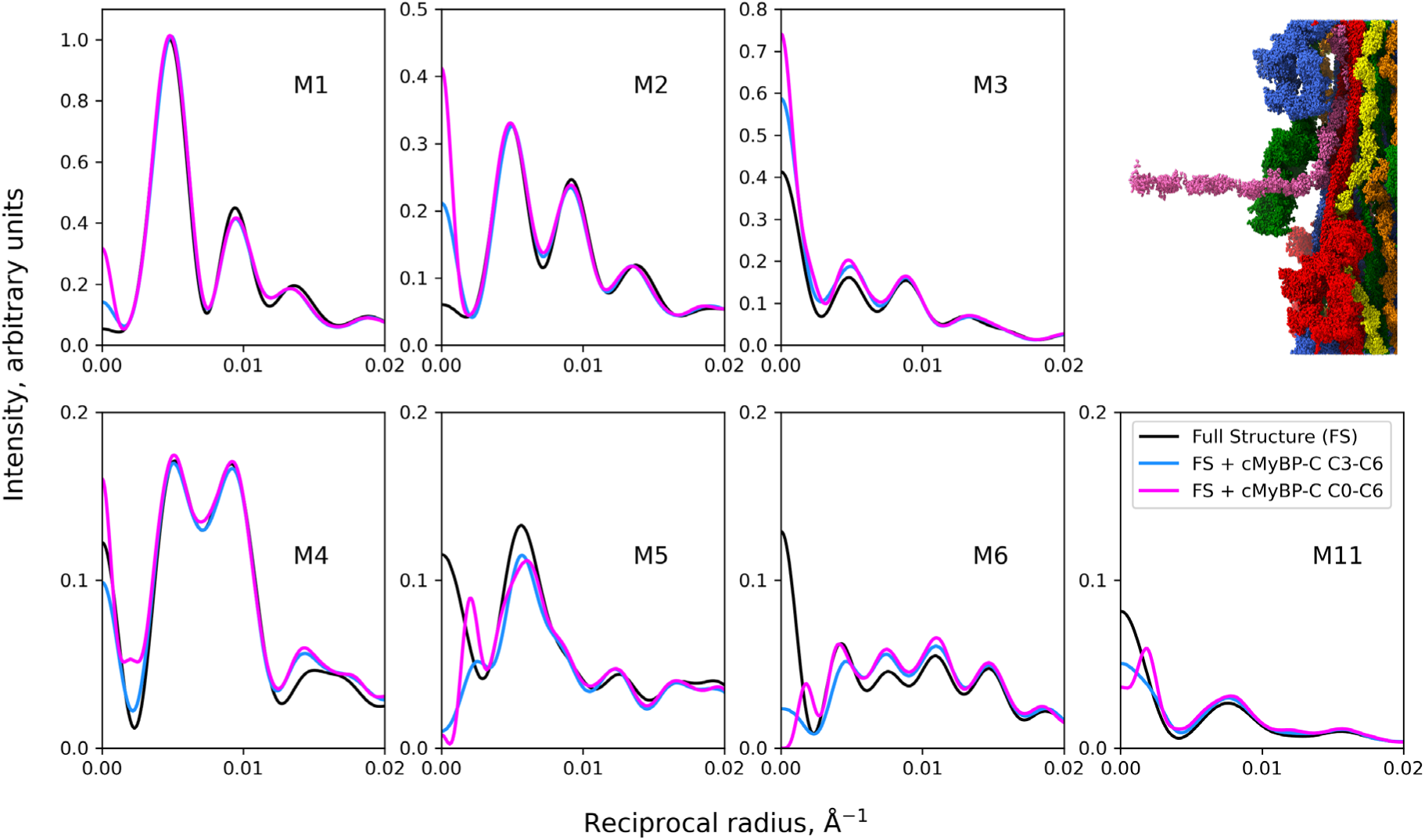
Computed intensities of myosin layer lines 1-6 and 11 from cardiac thick filament based on cryo-EM structure PDB 8g4l showing effect on meridian of extending domains C0-C6 of cMyBP-C radially from the thick filament. Black, full structure (8g4l), with domains C5-C10 extending along the surface of the thick filament; magenta, full structure but with domains C0-C6 extending at right angles to the filament (42,48) (top right); blue, same as magenta, but with domains C0-C2 removed, simulating their disorder. Radial extension of C0-C6 or C3-C6 has strong effects on all meridional reflections, either enhancing (M1-M3) or diminishing them (M5, M6; magenta, blue *vs.* black). The reduced meridional intensity on M5 and M6 is likely due to the orientation of C0-C6 exactly perpendicular to the filament axis, which produces strong peaks at higher orders of 430 Å and may have an amplitude at M6 similar in magnitude but opposite in sign to that of myosin heads, resulting in low intensity. However, with the broad distribution of angles of the C0-C6 fragment reported by (42), its contribution to the higher orders of 430 Å will be minimal, contributing to, and increasing the intensity of, the low-order reflections (M1-M3), but not to M6 and beyond.

It is important to note that in the sarcomere the N-terminal domains of cMyBP-C bind to actin, which has a periodicity of approximately 360 Å, rather than the 430 Å periodicity of myosin. This binding suggests a significant departure of C0-C6 domains from a perpendicular orientation to the filament axis in the actin-myosin lattice, consistent with cryo-EM data (42,50). Therefore, our calculations assuming perpendicular N-terminal domains, as shown in **Fig. 8**, likely give the maximum enhancement of meridional intensities resulting from the binding of these domains to actin.

### Contribution of titin and tails to forbidden meridionals

Myosin tails and titin also exhibit an axial repeat of 430 Å and, like cMyBP-C, are not helically organized. Although these components have little impact on the stronger region of the meridional pattern (M1-6), owing to their extended arrangement in the filament, they may significantly impact some of the weaker, higher order forbidden meridionals (M10, M11; (21)) (**Figs. 4, 5**).

In summary, the forbidden meridional reflections most likely arise from a combination of non-helically organized heads, partial disorder of CrD, and diffraction from titin and cMyBP-C (in the radially projecting configuration), with no component dominating, except for titin on M11.

## DISCUSSION

Interpretation of the X-ray diffraction pattern of vertebrate striated muscle has been hindered in past studies by the absence of a thick filament atomic model. We have used the model PDB 8g4l (23), based on cryo-EM of the human cardiac thick filament C-zone, to approximate the main coherently diffracting region of each half thick filament; contributions from the D-zone, P-zone and bare zone to the layer line intensities are likely to be minimal (see **Results: Atomic model of the C-zone used in computations**). Using this model, we elucidated the contributions of myosin, titin and cMyBP-C to the pattern; similar modeling was used to assess the contributions of the free and blocked heads of the IHM to the tarantula muscle X-ray diffraction pattern (51). We find that all components influence the pattern, with no single 1-to-1 connection between a particular constituent and reflection, with the exception of the dominant role of the major, bulky, globular components (myosin heads in the IHM configuration) in the strong, low angle region of the pattern (M1-8), and titin’s major contribution to the M11 meridional (see also **Supplemental Discussion: Effect of combining contributions from more than one component**). Elements extended along the filament axis (myosin tails, cMyBP-C, titin) contribute little to the strong reflections, but influence the weak, high angle region (M9-15), where the heads also contribute to a smaller extent. Partially disordered CrD heads and cMyBP-C radial extensions from the thick filament appear likely to be the main source of the forbidden meridional reflections.

### Explaining the forbidden meridional reflections

The origin of the forbidden meridional reflections in the vertebrate muscle X-ray pattern (those indexing on 430 Å rather 143 Å) has long been a puzzle (6), but must reflect components with a 430 Å projected density repeat (3,6,13,22). The strongest reflections are the M1 and M2 meridionals (**Fig. 1**). Our results (**Figs. 7, 8**) suggest that some disordering of CrD heads, possibly having a role as swaying (52)/‘sentinel’ heads (53), and/or cMyBP-C extensions towards actin filaments at 430 Å intervals are the main source of these strong reflections.

When skeletal muscle is stretched, decreasing overlap of the thin filaments with the thick filament C-zone, the M1 and M2 reflections become weaker (13). This supports the contribution to these reflections of cMyBP-C’s 430 Å-spaced radial extensions (domains C0-C6 (42)) bound to actin in the region of thin filament/C-zone overlap (13) (**Fig. 8**). At longer sarcomere lengths, availability of actin filaments for cMyBP-C binding would be reduced, leading to disordering of the N-terminal half of molecules nearer to the bare zone or possibly to the binding of these N-terminal halves along the thick filament (15). In either case, the consequent reduction in periodic fluctuation in protein density would lead to a drop in the meridional intensities (13), as observed. This would also explain reduction of the M2 meridional intensity in skeletal muscle when MyBP-C is cleaved between the C7 and C8 domains (54). The projecting C0-C7 domains would lose their 430 Å thick filament repeat and no longer contribute to the myosin X-ray reflections, including the forbidden meridionals.

The degree to which CrD heads are disordered, and how much this may also contribute to the strong (M1, M2) and other forbidden meridional reflections (**Fig. 7**), is unknown. In our reconstruction of isolated thick filaments, disordering is suggested by the noisy, weaker density of IHMs in crown CrD (23). But these IHMs on average still occupy their quasi-helical positions and have substantial density in the cryo-EM map (Fig. 1b of (23)), suggesting that disorder is limited. If so, this would support the strong forbidden meridionals (**Fig. 1**) coming primarily from cMyBP-C extensions.

### Origin of the M6 meridional

Changes in the M6 meridional spacing and intensity have been used to estimate thick filament compliance (55–57) and to model the crossbridge power stroke in contracting muscle (17–19). The absence of change in M6 intensity upon quick release of an isometrically contracting muscle, compared with a large change in M3 intensity, has been taken to mean that myosin heads in contracting muscle contribute little to M6. It has been suggested that this reflection arises, instead, from the myosin tails (17) or from the filament backbone as a whole, which would include cMyBP-C and titin as well as tails (18,19,38,56). Our results question this assumption, showing only a small contribution of these backbone components to M6 and other meridionals, except for M11 (**Figs. 4-6**), consistent with their extended structure and minor longitudinal periodic mass variation. Our modeling suggests a significant contribution of the myosin heads to the M6 meridional intensity in relaxed muscle (**Fig. 3**), supported by a notable decrease in intensity upon a reduction of ordered myosin heads by various perturbations (58,59). Employing the M6 meridional reflection as a measure of changes in the length of myosin filaments may therefore not be justified. In contrast, the meridional intensity on M11 originates mainly from titin (**Fig. 5**; (23)), which binds to the myosin tails (1,23), and has little contribution from the heads (**Fig. 3**). This makes the spacing of this reflection suitable for estimating elongation or shortening of the filaments. The spacing of M11 increased by 1.56-1.87% during transition of relaxed frog muscles to isometric tetanic contraction and increased further by 0.16% upon stretch of contracting muscles (57).

During the transition of relaxed skeletal muscle fibers to isometric contraction, the M6 meridional intensity increases by a factor of 1.68 (38). Since the contribution of tails, titin and other components associated with the backbone of the thick filaments seems unlikely to change, this suggests a major contribution of the myosin heads to M6 intensity in contracting muscle. This can be explained by the fact that a large fraction of myosin heads binds actin, while maintaining myosin backbone periodicity. Those nearly perpendicular to the filament axis would be expected to contribute strongly to the M3 and M6 meridional intensities.

The question remains: if the M3 and M6 reflections both have substantial contributions from myosin heads, why do their intensities and spacings not behave in lockstep? For example, it has been shown that M3 and M6 spacings show different time courses during force generation (60,61) and different behavior in response to various perturbations such as passive stretch (13,60) or differences in the proportions myosin heads in the ordered state (59,62) in different muscle types. It seems, therefore, that the M3 and M6 reflections are not reporting the same underlying phenomena. An ultimate explanation of the different behaviors of the M3 and M6 requires more complex models than those considered here, incorporating information from high-resolution structures of the P- and D-zones, and also accounting for the 2D actin-myosin lattice and the different populations of strongly and weakly bound actin-myosin complexes in contracting muscle (17,63). Such models lie outside the scope of this paper and should be addressed in the future.

### How well does the reconstruction fit the experimental X-ray pattern?

In the analysis presented here, we have used PDB 8g4l, an atomic model of the C-zone repeat of the cardiac thick filament (23), to estimate how different thick filament components contribute to the myosin layer line pattern of relaxed cardiac muscle. But how closely does the predicted thick filament pattern match experimental data (**Fig. 1**)? To answer this, we ignore the thin filaments, which, in relaxed muscle, diffract independently of the thick filaments and give rise to layer lines indexing on the ∼360 Å helical repeat of actin (**Fig. 1: A6, A7**) (6). Most myosin layer lines (based on a 430 Å repeat) are separate from the actin layer lines, and can be extracted separately (**Fig. 1b**), except M7 and M8, which overlap with the strong A6 and A7 actin layer lines. We have therefore compared our calculated thick filament pattern, based on the C-zone atomic model (8g4l), with the observed diffraction on myosin layer lines M1-M11, except M7 and M8 (**Fig. 9**).

**Fig. 9.**
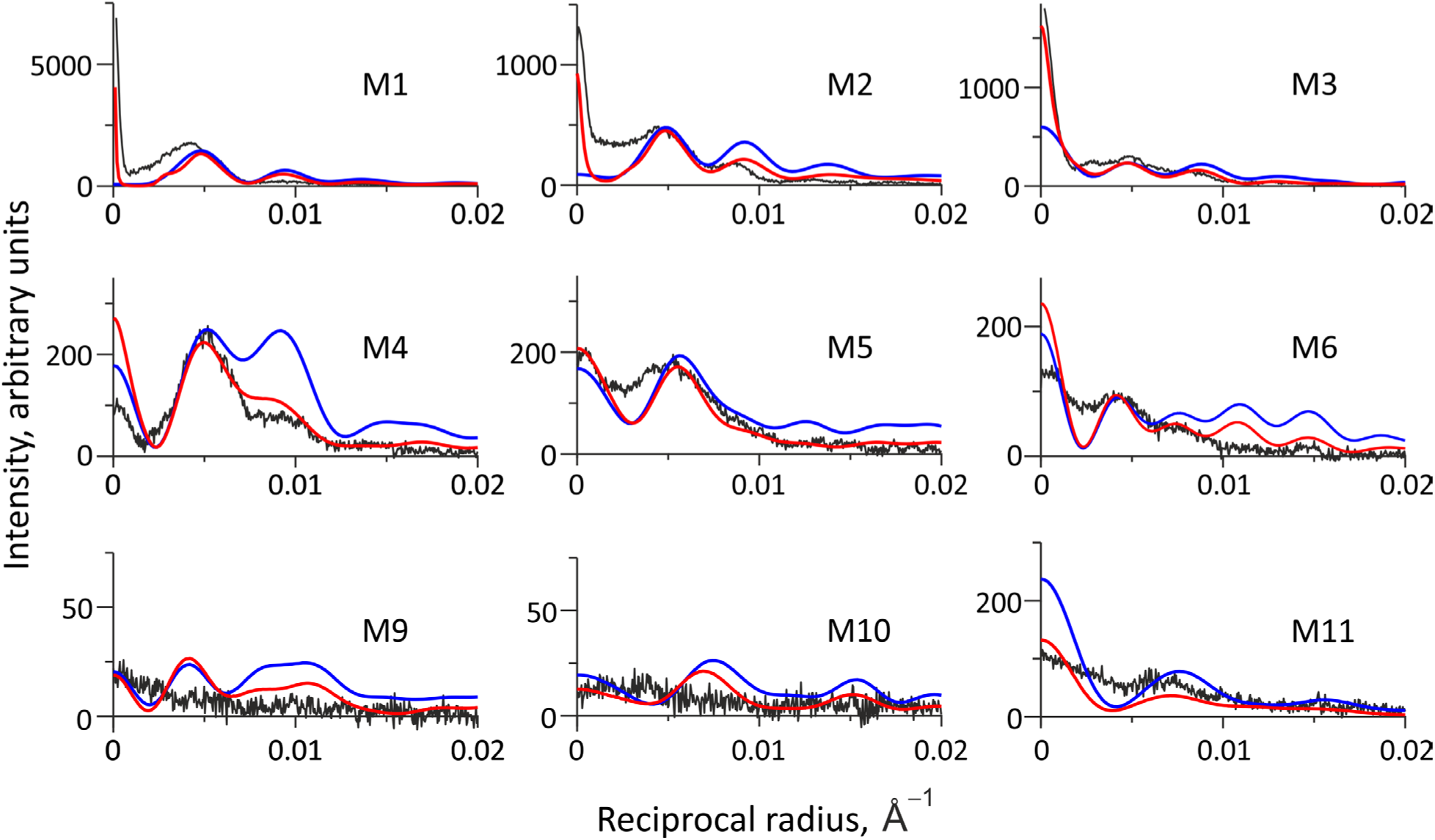
Comparison of experimental X-ray pattern with computed myosin layer line intensities based on cardiac thick filament atomic model (PDB 8g4l). Black, background-subtracted experimental layer line intensities (Fig. 1b). Blue, computed intensities for the full atomic structure (PDB 8g4l), representing a single filament. Red, computed intensities for the same model placed within a hexagonal filament lattice, incorporating disorders of the first and second kind (36): (1) Disorder of the first kind: rotational and radial “thermal” disorder of the myosin heads (standard deviations of 10° and 10 Å, respectively, relative to their model positions; *cf.* Figs. S3, S4); (2) Disorder of the second kind: radial and axial disorder in filament packing within the lattice (standard deviations of 100 Å radially and 45 Å axially; **Fig. S2**). Positioning filaments within the lattice greatly increases intensity of the meridional peaks on layer lines M1-M3 (red vs. blue) and narrows them due to the longitudinal registration of neighboring filaments (see text), helping to explain the strong meridional reflections in the experimental pattern. The “temperature factor” in (1), reflecting the thermal mobility of myosin heads in a real filament compared to their rigid structure in the atomic model, accounts for the intensity fall-off at higher reciprocal radii, including the weakening of the second peak on the M4 layer line to the level seen in experimental patterns (red vs. blue). The high intensity of the M11 meridional in the model pattern (blue) is reduced close to the experimental level (black) with a ∼7.5 Å axial mobility of titin domains (**Fig. S5**). The intensities were scaled using a factor that provides a fit of the higher angle half of the strong off-meridional peak on the M1 layer line in the calculated and experimental patterns. This approach accounts for the possible contribution of disordered myosin heads – primarily in the D-zone – to the near-meridional intensity on M1. This scaling choice also resulted in a reasonably good fit for the other myosin layer lines.

The myosin layer lines in the experimental pattern show no obvious sampling by the hexagonal thick filament lattice (there is no evidence for a simple or super lattice (64)), implying that the filaments have random rotations about their long axis and/or that there is significant variation in the distances between neighboring filaments. In the absence of sampling, each myosin layer line approximates the transform profile of a single thick filament, as we have used for our modeling, validating this method of comparison with experimental data (**Fig. 9**). The fit of 8g4l (blue) to the experimental pattern (black) was reasonable in the stronger part of the pattern (**Fig. 9**, M1-M6), with the exception of the meridian, which is underestimated in the model, especially on M1-M3, and overestimated on M11 (blue *vs.* black traces). At higher reciprocal radii, the pattern from the model is stronger than seen experimentally (**Fig. 9**, blue *vs.* black), including the second peak on the M4 layer line, predicted by the model, but weak in the data. These discrepancies have straightforward explanations, as detailed below.

Meridional reflections in experimental patterns are affected not only by the structure of the single unit cell that we have been modeling (one 430 Å repeat of the C-zone), but also by incorporation of thick filaments in longitudinal register into the hexagonal lattice of the sarcomere (11,17). The experimentally observed high intensity and narrow peaks of the M1-M3 meridionals (**Fig. 9**, black) suggest strong axial registration of thick filaments, consistent with EM observation of sectioned cardiac muscle (35,49). When we introduce such lattice registration into our model, the M1-M3 meridional intensities are strongly intensified (**Fig. S2**, **Fig. 9**, red *vs.* blue) and better match the observed intensities (**Fig. 9**, red *vs.* black); this is true whether cMyBP-C is extended along the filament as in 8g4l (**Fig. 9**, red) or the N-terminal half extends radially (**Fig. S2**, orange).

The stronger intensities predicted by the model at high reciprocal radii (**Fig. 9**, blue) compared to the experimental data (black) are likely to be due to thermal motion of the myosin heads in an intact muscle compared with their static, fixed positions in the atomic model. This can be simulated by incorporating “thermal” disorder (or disorder of the first kind (36)) of the myosin heads into the predicted pattern. This disorder includes both rotational (azimuthal) and radial deviations of the IHM positions from their location in the cryo-EM-derived model of the 430 Å-long C-zone repeat. Assuming that both components follow Gaussian distributions, with realistic standard deviations of 10° and 10 Å, respectively, the computed layer lines weaken at higher reciprocal radii (**Figs. S3, S4**), bringing them into close agreement with the experimental pattern (**Fig. 9**, red *vs.* black). The rotational disorder effectively reduces the contribution of the high-order Bessel functions to the diffraction intensity (such as the second peak on the M4 layer line), while the radial disorder decreases the intensity at high reciprocal radii (see Formula SM4 in Supplementary Methods, Supporting Materials and red lines in **Fig. 9**). Similarly, the weaker measured intensity of the M11 meridional reflection, coming mainly from titin, compared with the model prediction (black *vs.* blue, **Fig. 9**) can be explained by small axial motions of titin domains on the filament surface (**Fig. S5**). When accounted for, these motions predict an M11 intensity similar to that observed (**Fig. 9**, M11, red *vs.* black).

## CONCLUSION

Our analysis of the X-ray diffraction pattern produced by the vertebrate thick filament, and the contributions of different components in the C-zone to specific layer lines, demonstrates that the C-zone is the main contributor to the diffraction pattern of muscle in the relaxed state at physiological temperature. This provides a quantitative foundation for future X-ray studies of the contractile mechanism, as well as the impact of disease mutations and therapeutic drugs on muscle molecular structure. While ultimately we would like to explain the experimental pattern in full, this requires more detailed knowledge of thick filament structure, which should come from further EM and other biophysical studies. The structure of the C-zone of isolated thick filaments from human cardiac muscle (23) is similar to that obtained by cryo-electron tomography (cryo-ET) of mouse cardiac thick filaments in the intact, relaxed myofibrillar lattice (42). The main difference is the radial projection of cMyBP-C N-terminal domains towards actin in the myofibril, which helps to explain the forbidden meridional reflections, as discussed above. The similarity of our isolated filament structure to that in the intact myofibril supports its validity for annotating the X-ray pattern of intact muscle. Cardiac and skeletal muscles have similar X-ray patterns (7,12,60,65) and similar thick filament EM images (22,66), suggesting that their thick filaments have similar structures and that the annotations we have made are also relevant to skeletal muscle.

## Author Contributions

NAK designed research, performed research and analyzed data; DD performed research; WM performed research; AKT designed research, performed research and analyzed data; TI analyzed data; RP designed research and analyzed data; RC designed research, analyzed data and wrote the paper, with input from NAK, AKT and RP.

## Acknowledgments

We thank Dr. Maicon Landim-Vieira, Mr. Vivek Jani and Mr. Christopher McAllister for help with the X-ray experiments. This work was supported by NIH grants AR072036, HL139883 to RC, HL164560, and AR081941 to RP, and was conducted under the state assignment of Lomonosov Moscow University (NK). UCSF ChimeraX, used for molecular graphics in Fig. 2, was developed by the Resource for Biocomputing, Visualization and Informatics at the University of California, San Francisco, with support from NIH grant GM129325 and the Office of Cyber Infrastructure and Computational Biology, National Institute of Allergy and Infectious Diseases. X-ray diffraction experiments were supported by grant P30 GM138395 from the National Institute of General Medical Sciences of the National Institutes of Health and were conducted at the Center for High-Energy X-ray Sciences (CHEXS), supported by the National Science Foundation (BIO, ENG and MPS Directorates) under award DMR-1829070, and at the Macromolecular Diffraction at CHESS (MacCHESS) facility, supported by award 1-P30-GM124166-01A1 from the National Institute of General Medical Sciences, National Institutes of Health, and by New York State’s Empire State Development Corporation (NYSTAR).

## Data-sharing Plans

Data will be deposited in the Open Science Framework public repository. This includes original X-ray diffraction patterns and extracted layer-lines.

## Supplemental Information

**Fig. S1.**
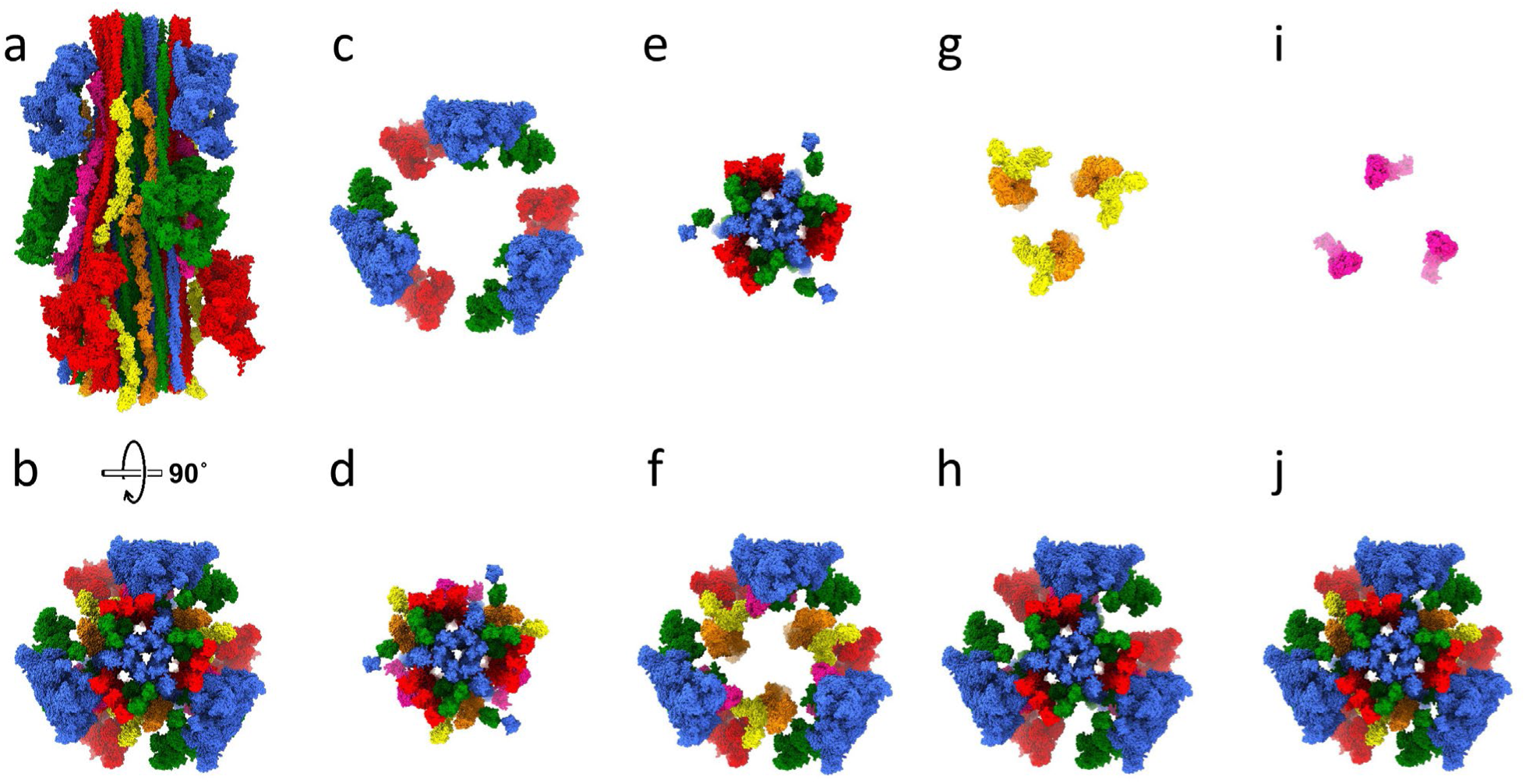
Human cardiac thick filament atomic model of 430 Å repeat of C-zone (PDB: 8g4l) showing the individual components alone or removed, in transverse view; compare with. **Fig. 2b-k. (a)** 430 Å C-zone repeat with all components, longitudinal view. Red, green and blue, CrD, CrH and CrT IHMs, respectively; orange and yellow, titin A and B, respectively; pink, cMyBP-C domains C5-C10. **(b)** Full structure viewed transversely towards Z-line. **(c, d)** Same as **(b)** but with heads only **(c)** or heads removed **(d)**. **(e, f)** Same as **(b)** but with tails only **(e)** or tails removed **(f)**. **(g, h)** Same as **(b)** but with titin only **(g)** or titin removed **(h)**. **(i, j)** Same as **(b)** but with cMyBP-C only **(i)** or cMyBP-C removed **(j)**.

**Fig. S2.**
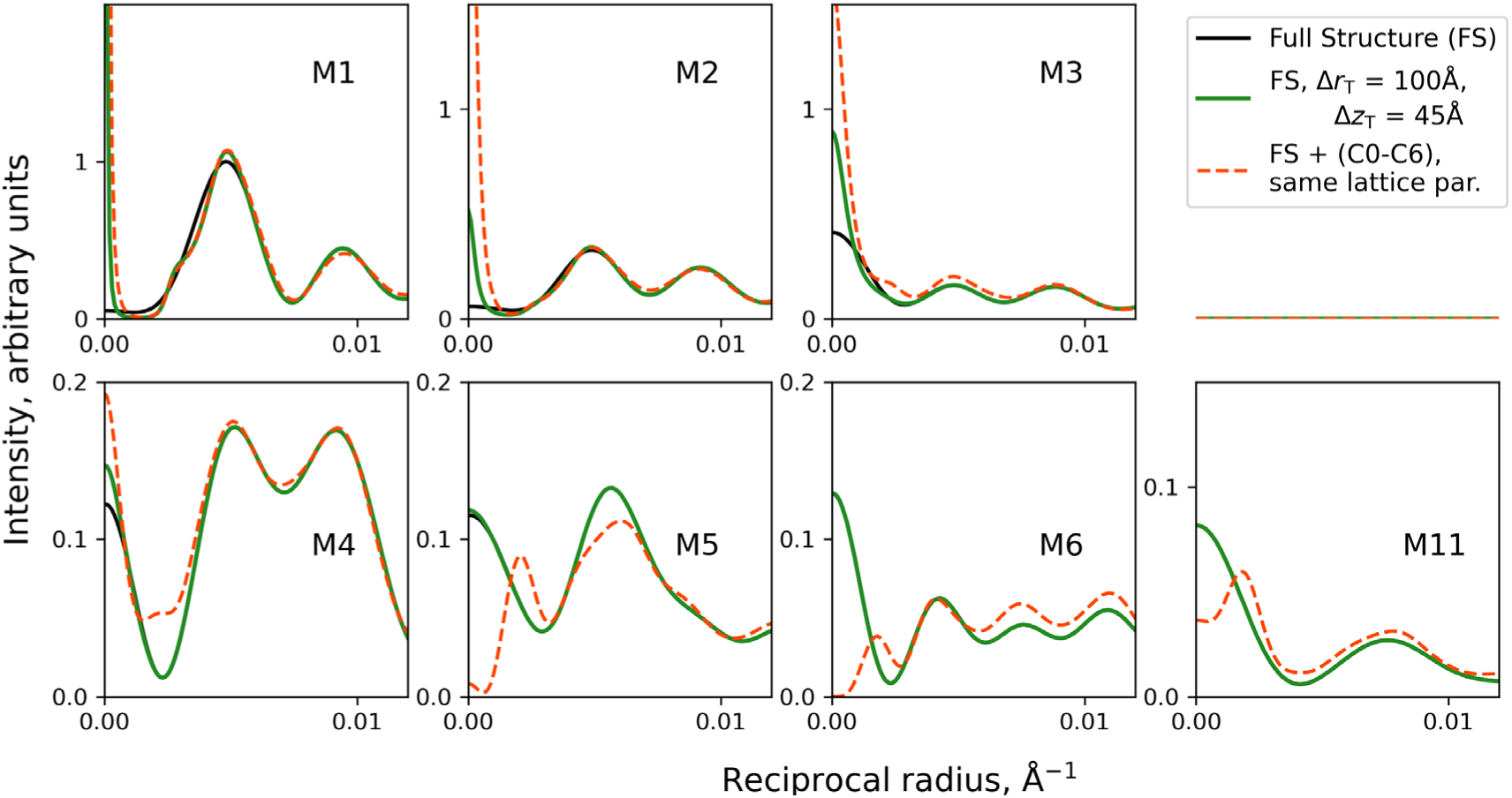
Placing filaments in hexagonal lattice enhances low order meridional intensities. Black, full structure (PDB 8g4l) of single filament (no lattice), as in **Figs. 3-8**, black traces. Green, same as black, but with filaments placed in hexagonal lattice, in close longitudinal register (Δz_T_ = 45 Å) and with radial (azimuthal) disorder of filament positions of Δr_T_ = 100 Å. Orange, same as green, but with cMyBP-C domains C0-C6 extended radially (as in Fig. 8) (42,48). Longitudinal registration causes dramatic intensification of meridional intensity on M1 and M2 (green *vs.* black), as observed experimentally—even when cMyBP-C runs longitudinally, as in 8g4l. There is further intensification (M1-M4) when C0-C6 is perpendicular—orange here (in lattice) *cf.* Fig. 8 (magenta, no lattice). Δr_T_ chosen so that there is no lattice sampling on off-meridional parts of layer lines, as observed experimentally (see text); thus, off-meridional regions of black trace lie immediately under green trace. Δr_T_ = 100 Å and Δz_T_ = 45 Å are used in Fig. 9, red trace.

**Fig. S3.**
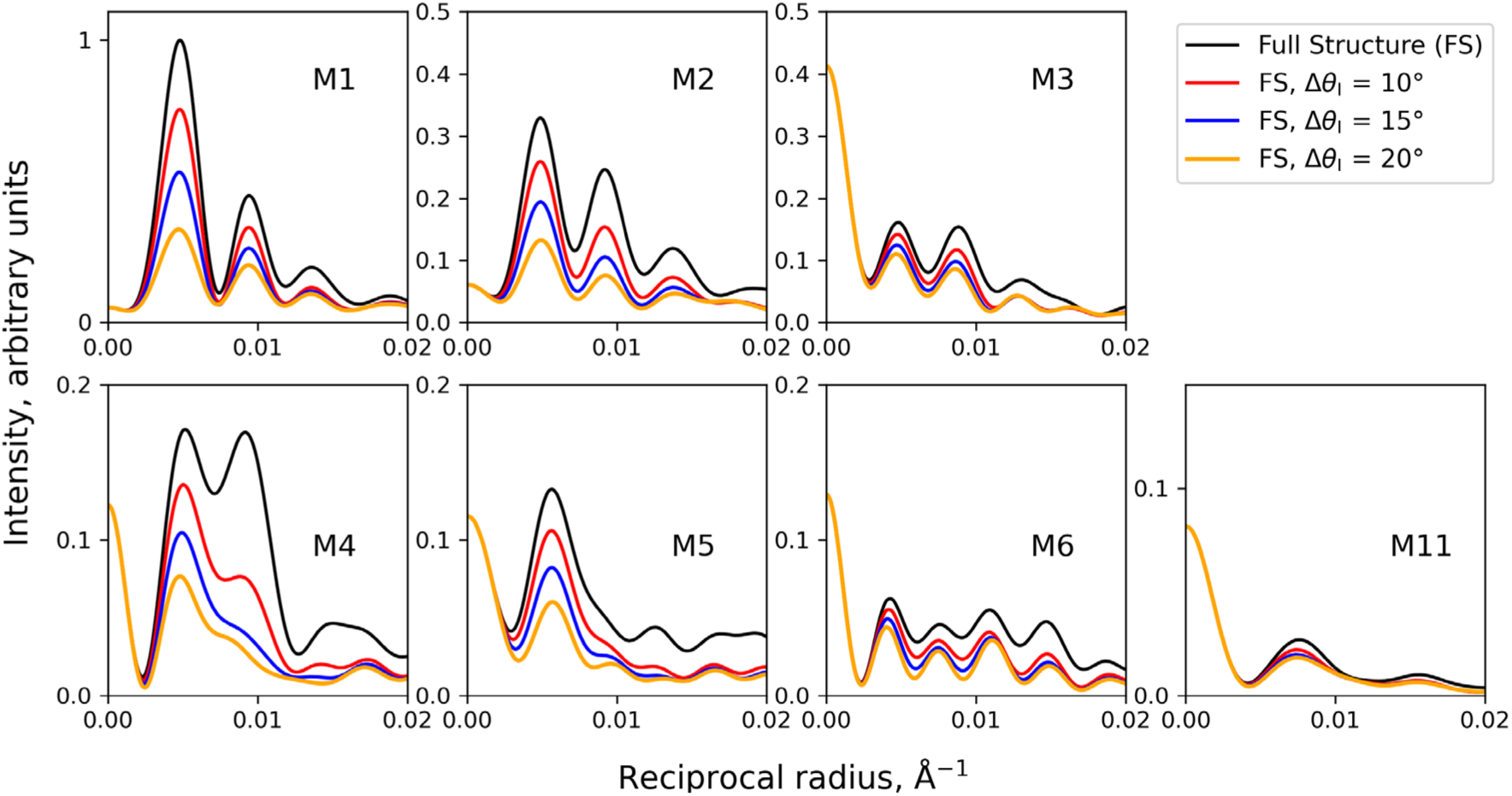
Rotational (azimuthal) disordering of IHMs reduces the contribution of high-order Bessel functions to the diffraction intensity. The traces show the predicted layer line intensities from a single filament (FS) as in **Figs. 3-8** (black), and the same structure with heads azimuthally disordered by different amounts, Δθ_I_ (red, blue, orange). The traces coincide near the meridian, but become weaker at higher reciprocal radii as disorder increases. There is a dramatic reduction of the second peak on the M4 layer line. Δθ_I_ = 10° is used in Fig. 9, red trace.

**Fig. S4.**
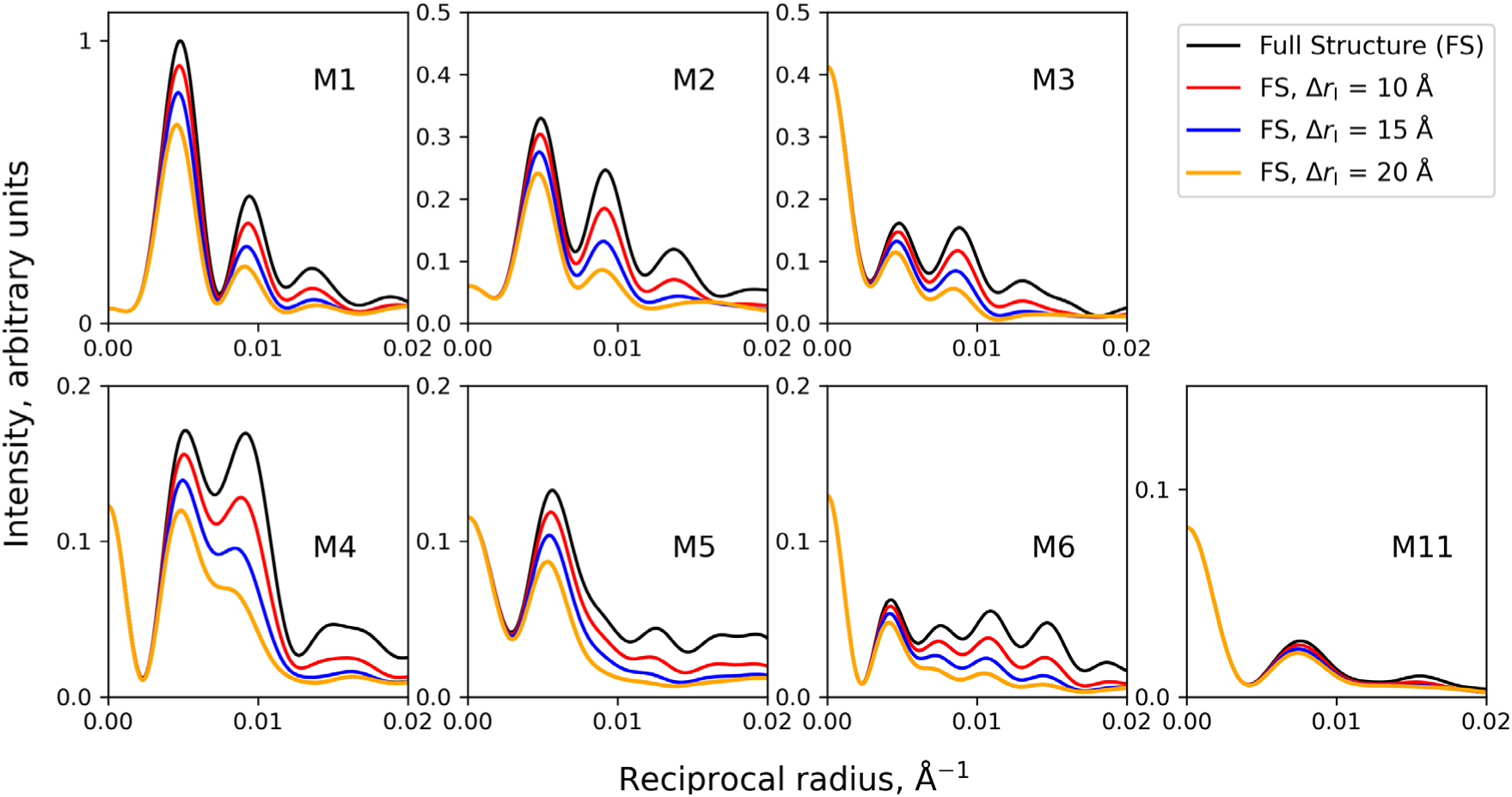
Radial disordering of IHMs decreases off-meridional intensities especially at higher reciprocal radii. The traces show the predicted layer line intensities from a single filament (FS) as in **Figs. 3-8** (black), and the same structure with heads radially disordered by different amounts, Δr_I_ (red, blue, orange). The traces coincide near the meridian, but become weaker at higher reciprocal radii as radial disorder increases. Δr_I_ = 10 Å is used in Fig. 9, red trace.

**Fig. S5.**
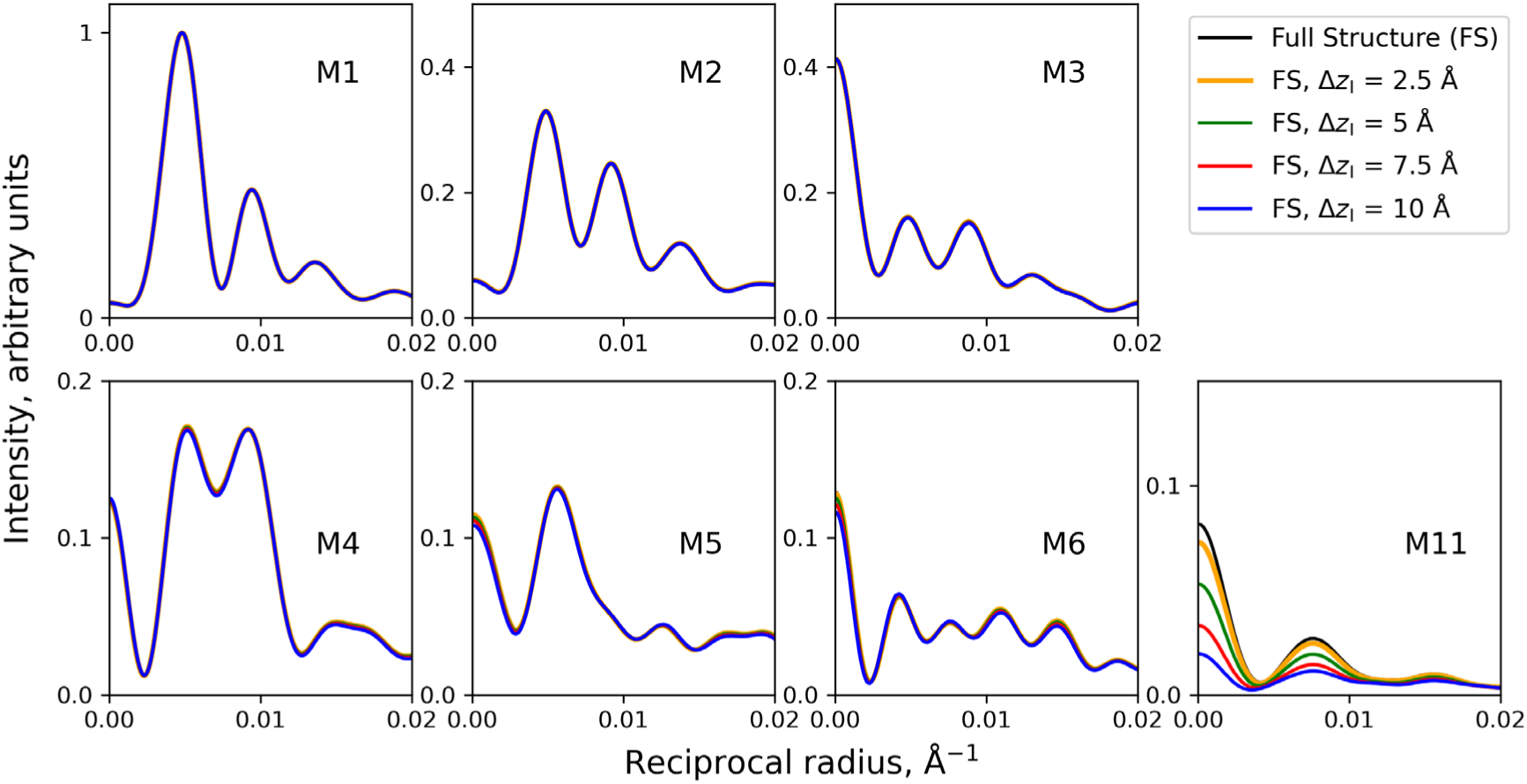
Longitudinal disordering of titin domains decreases M11 intensity, with no effect on other layer lines. The traces show the predicted layer line intensities from a single filament (FS) as in **Figs. 3-8** (black), and the same structure with titin domains longitudinally disordered by different amounts, Δz_I_ (orange, green, red, blue). M11 becomes weaker with increased disorder. Δz_I_ = 7.5 Å is used in Fig. 9, red trace.

## Supplemental Methods

### Calculation of layer line intensities

Calculations were performed using cylindrical coordinates (*r, 𝑟 z*) with the *z*-axis coinciding with the thick filament axis. The axial size of the unit cell was equal to the axial period of the original atomic structure, 8g4l, *c* = 430 Å. The total number of atoms in the model (see inset in **Fig. 3**) was 377,364. Each heavy (non-hydrogen) atom in the unit cell was assigned three coordinates (*r_j_, 𝜓_j_, 𝑧_j_*) where *j* represents the atom number in the model. Fourier transforms 𝐹𝐹_𝑙𝑙_ on the *l*-th myosin layer line were calculated assuming an infinite number of axial repeats of the unit cell according to the formula (36):

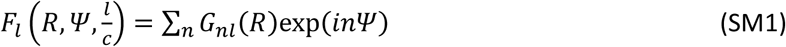

where 𝑅𝑅, 𝛹𝛹 are the radial and azimuthal coordinates in reciprocal space. 𝐺𝐺_𝑛𝑛𝑙𝑙_ is the Fourier-Bessel structure factor:

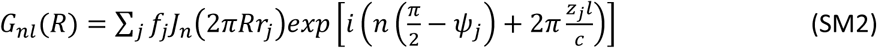

where 𝑓𝑓_𝑗𝑗_ is the scattering factor of the *j*-th atom, equal to the number of electrons in the atom minus the electron density of water within a sphere corresponding to the Van-der-Waals radius of the atom (41); 𝐽𝐽_𝑛𝑛_ is the *n-*th order Bessel function of the first kind. Due to 3-fold symmetry of the original data, the total number of atoms used for calculations was reduced: 377,364/3 = 125,788, and only 𝐽𝐽_𝑛𝑛_ with *n* multiples of 3 were counted.

### Incorporating disorder into computations

Assume that the IHMs in the structure behave as rigid bodies that are subject to “thermal” disorder (disorder of the 1st kind) (36), with both angular and radial deviations following Gaussian distributions characterized by root-mean-square deviations Δ𝜃𝜃_I_ and Δ𝑟𝑟_I_, respectively. We also assume that titin is subject to axial “thermal” disorder with root-mean square deviation Δz_I_. Under these assumptions, expressions (SM1)-(SM2) can be rewritten as (67):

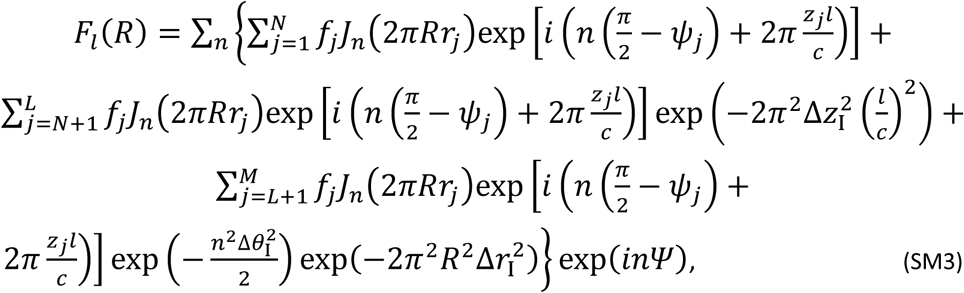

where atoms numbered 𝑗𝑗 = 1, …, 𝑁𝑁 correspond to the myosin tails and MyBP-C, while atoms numbered 𝑗𝑗 = 𝑁𝑁 + 1, …, 𝐿𝐿 correspond to titin, and those with numbers from 𝐿𝐿 + 1 to 𝑀𝑀 represent the myosin heads subjected to “thermal” disorder.

The intensity of the layer lines was then calculated as the azimuthally averaged square of the Fourier transform, assuming infinite lattice disorder:

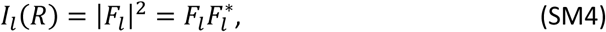

where * here and hereafter means complex conjugate.

This classical approach allowed us to estimate the impact of all protein components on the off-meridional intensities of the myosin layer lines. However, the observed intensities of the meridional reflections were higher than those simulated in an infinite disorder model.

Alternatively, the unit cell arrangement, including axial (*z* axis) and radial (*x, y* plane) disorder, was incorporated into the calculations based on the translation of myosin filaments within the lattice, as previously suggested (28,36). To the best of our knowledge, no solid experimental data is available on the lattice organization of myosin filaments in mammalian heart muscle sarcomeres. For simplicity, we assumed a simple hexagonal lattice (64). No indication of lattice sampling was observed in the diffraction patterns except on the meridian (**Fig. 1**), indicating a high degree of radial filament disorder. Consequently, there is no practical difference between assuming a simple or a super-lattice (64) when calculating off-meridional layer line intensities. In this context, the layer line intensity was calculated as follows. The interference function, 𝑍𝑍, which accounts for the axial and radial translation of filaments in the hexagonal lattice (neglecting filament tilt and rotation) under a disorder of the second kind, is expressed as (28,36,68):

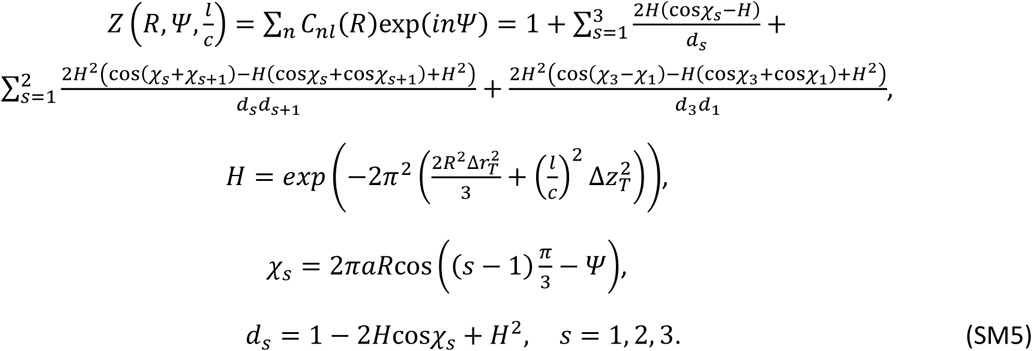

Here, 𝐶𝐶_𝑛𝑛𝑙𝑙_ are the Fourier series coefficients of the interference function, 𝑍𝑍, Δ𝑟𝑟_𝑇𝑇_ and Δ𝑧𝑧_𝑇𝑇_ represent the root-mean square deviations of the radial and axial positions of the myosin filaments in the lattice, and *a* denotes the length of the vector of transversal translation of the unit cells in the lattice. For a simple lattice, *a* equals the distance between neighbor myosin filaments corresponding to the unit cell size. The disorder is assumed to be of the second kind, or paracrystalline type, as neighboring myosin filaments are connected at the M-line in the center of the sarcomere by myomesin, C-termini of titin, obscurin, and possibly other proteins.

The layer line intensity was calculated as

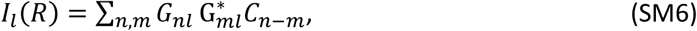

where G^*∗^ is the complex conjugate of 𝐺𝐺_𝑚𝑚𝑙𝑙_.

To simulate the experimental diffraction pattern obtained in the presence of mavacamten (**Fig. 9**), we selected the following disorder parameters: Δ𝜃𝜃_I_ = 10° and Δ𝑟𝑟_I_ = 10 Å for the “thermal” azimuthal and radial IHM disorder, and Δ𝑧𝑧_I_ = 7.5 Å for the axial “thermal” disorder of titin. The latter value is comparable to the resolution of cryo-EM data. For lattice disorder we chose Δ𝑟𝑟_𝑇𝑇_ = 100 Å, Δ𝑧𝑧_𝑇𝑇_ = 45 Å. The interfilament distance, 𝜋𝜋 = 400 Å, was determined from the position of the 1.0 equatorial reflection, while Δ𝑧𝑧_𝑇𝑇_ was estimated from the radial width of the M1 meridional reflection. The high value of Δ𝑟𝑟_𝑇𝑇_ leads to the absence of visible lattice sampling on the off-meridional layer lines, consistent with experimental observations.

## Supplemental Discussion

### Effect of combining contributions from more than one component

To simplify analysis, we studied the impact of each individual thick filament component on the different layer lines. While myosin tails, titin and cMyBP-C have only small individual effects (**Figs. 4-6**), what happens when they are combined, as in the native filament? When all 3 components are removed together, leaving only the heads (**Fig. 3**, red vs. black), there is relatively little impact on the stronger layer lines (M1-5) when compared with removal of tails only, but a substantial impact on M6, 8, 10, 11, 12 and 15 (**Fig. 4**, blue). Removal of the three components (**Fig. 3**, red), compared with removal of titin only (**Fig. 5**, blue), shows little impact, except for minor effects on M6, 9 and 14. Removal of the three components (**Fig. 3**, red) compared with removal of cMyBP-C only (**Fig. 6**, blue), also has only a small impact, with minor differences on M6, 8, 9, 10, 11, 15. In summary, while a combination of all three extended structures affects the pattern more than the individual components, the impact is generally small and mostly confined to the low intensity, higher order layer lines.

These results illustrate that the contributions of different components to the X-ray intensities are not additive. This is because intensity is the square of the sum of the complex scattering amplitudes from each contributing component (equal to the Fourier transform of the electron density): because the components have a fixed spatial relationship to each other (phase), they cannot be considered independently, but as coherently linked diffractors. Thus, while the scattered amplitudes from different components are additive, the intensities are not (36,69).

For example, heads alone produce roughly half the intensity on the M6 meridional reflection (**Fig. 3, M6, red**), while the balance of the structure (tails, titin and cMyBP-C, left after removing the heads; **Fig. 3**, M6, blue) produces 13%, not 50%. For M9 (**Fig. 3**), the intensities of heads alone (red) and the structure lacking heads (blue) are both greater than that for the full structure.

Like any complex value, the amplitude can be represented either as a sum of real and imaginary parts or by its magnitude and phase. For example, with the M6 meridional (**Fig. 3**), the Fourier transforms or amplitudes, 𝐹𝐹, for the full structure (𝐹𝐹_𝐹𝐹𝐹𝐹_, black), heads only (𝐹𝐹_𝐻𝐻_, red) and minus heads (𝐹𝐹_𝑀𝑀𝐻𝐻_, blue) at the meridian (i.e. at a reciprocal radius 𝑅𝑅 = 0) expressed in arbitrary units are approximately:

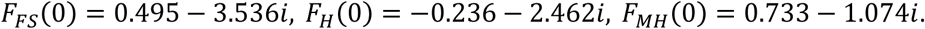

Thus, for amplitudes, 𝐹𝐹_𝐹𝐹𝐹𝐹_(0) = 𝐹𝐹_𝐻𝐻_(0) + 𝐹𝐹_𝑀𝑀𝐻𝐻_(0).

The intensity of each reflection, 𝐼𝐼, is the sum of the squares of the real and imaginary parts. At the M6 meridian we have:

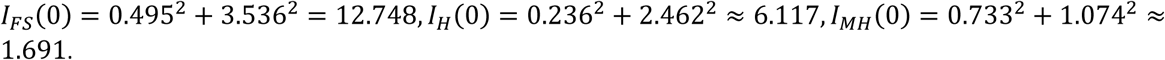

Thus, for intensities, 𝐼𝐼_𝐻𝐻_(0) + 𝐼𝐼_𝑀𝑀𝐻𝐻_(0) ≠ 𝐼𝐼_𝐹𝐹𝐹𝐹_(0).

Also, the magnitudes are not additive due to phase differences: |𝐹𝐹_𝐻𝐻_(0)| + |𝐹𝐹_𝑀𝑀𝐻𝐻_(0)| ≠ |𝐹𝐹_𝐹𝐹𝐹𝐹_(0)|

where

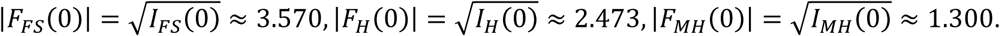

The heads contribute to 𝐼𝐼_𝐻𝐻_ (0)⁄𝐼𝐼_𝐹𝐹𝐹𝐹_(0) ≈6.117/12.748 ≈ 48% of the M6 intensity on the meridian—consistent with **Fig. 3** (M6, magenta).

Similar to the above, for titin’s contribution to the M11 meridional intensity (**Fig. 5**), we obtain:

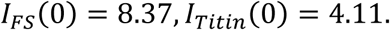

Thus the apparent titin contribution is 4.11/8.37 ≈ 49% of the total M11 intensity. However, when considering the magnitudes (absolute values) of the Fourier transforms the contribution differs:

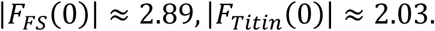

Thus titin contributes 2.03/2.89 ≈ 70% of the M11 magnitude. The other components (tails, heads and cMyBP-C) contribute 29% (0.85/2.89) of the magnitude, although their intensity is only 9% (0.73/8.37) of the full intensity (**Fig. 5**): 𝐼𝐼_𝑀𝑀𝑇𝑇_(0) ≈ 0.73, |𝐹𝐹_𝑀𝑀𝑇𝑇_(0)| ≈ 0.85, where MT refers to “minus titin”.

### Quantification of the impact of mavacamten on thick filament order

The C-zone atomic model that we used to calculate the layer line intensities was based on a cryo-EM structure derived from cardiac muscle thick filaments in the presence of the myosin inhibitor, mavacamten, used to maximize resolution (23) (similar to (42)). It therefore represents an “idealized” relaxed thick filament, similar to, but more stable than, a physiologically relaxed filament (70); for consistency, the experimental pattern we used for fitting (**Fig. 1a, b**) was also made with mavacamten-treated muscle. So, how closely do our annotations apply to physiologically relaxed muscle? While we cannot exclude the possibility that mavacamten might be artificially enriching the ordered populations away from physiologically realistic values, its stabilization of the IHM appears to occur without significant alteration of overall IHM structure. This is implied by the close similarity, at near-atomic resolution, of the isolated IHM (PDB 8act; (71)), where no mavacamten was used, to the two stable IHMs in our cryo-EM structure (crowns CrH and CrT of PDB 8g4l) (23). It is also suggested by the similarity, at lower resolution, of our reconstruction to that of cardiac thick filaments treated with blebbistatin (72) and without any drug (46), and to thick filaments in fish skeletal muscle, also without drugs ((11) *cf.* (23)).

To test this further, we also obtained diffraction patterns from skinned porcine cardiac muscle in the absence of mavacamten in parallel with our mavacamten patterns. Because these were of insufficient quality to extract the radial distribution profiles of diffraction intensity for individual myosin layer lines, we quantitatively evaluated the effect of mavacamten by analyzing the total off-meridional intensities within a reciprocal radial range of 0.0015–0.015 Å⁻¹. To ensure comparability across patterns from different samples, these intensities were normalized to the actin layer line A6 (at ∼59 Å), based on the assumption that actin filaments are unaffected by mavacamten. In the absence of mavacamten, the normalized off-meridional intensities of myosin layer lines M1–M6 were reduced by 14–29% compared to those in its presence (see figure below). For the M1 layer line specifically, the normalized intensity (M1/A6) was 1.91 with mavacamten and 1.66 without it. This 13% reduction in normalized intensity (27% reported previously (70)), reflects a 7% (or 11% (70)) reduction in amplitude of the M1 layer line in the absence of mavacamten, suggesting a very mild stabilization of myosin head helical order in its presence. This contrasts with cryo-ET of relaxed cardiac muscle without mavacamten, which shows mostly disordered myosin heads (73) and would therefore be expected to produce weak or non-existent layer lines, contrary to X-ray diffraction evidence (49,70). Disorder in this case may be due to phosphorylation of cMyBP-C or the regulatory light chains (73–75), or to preparative or other processes.

We conclude that our annotations are relevant to physiologically relaxed cardiac thick filaments, in the absence of drug.

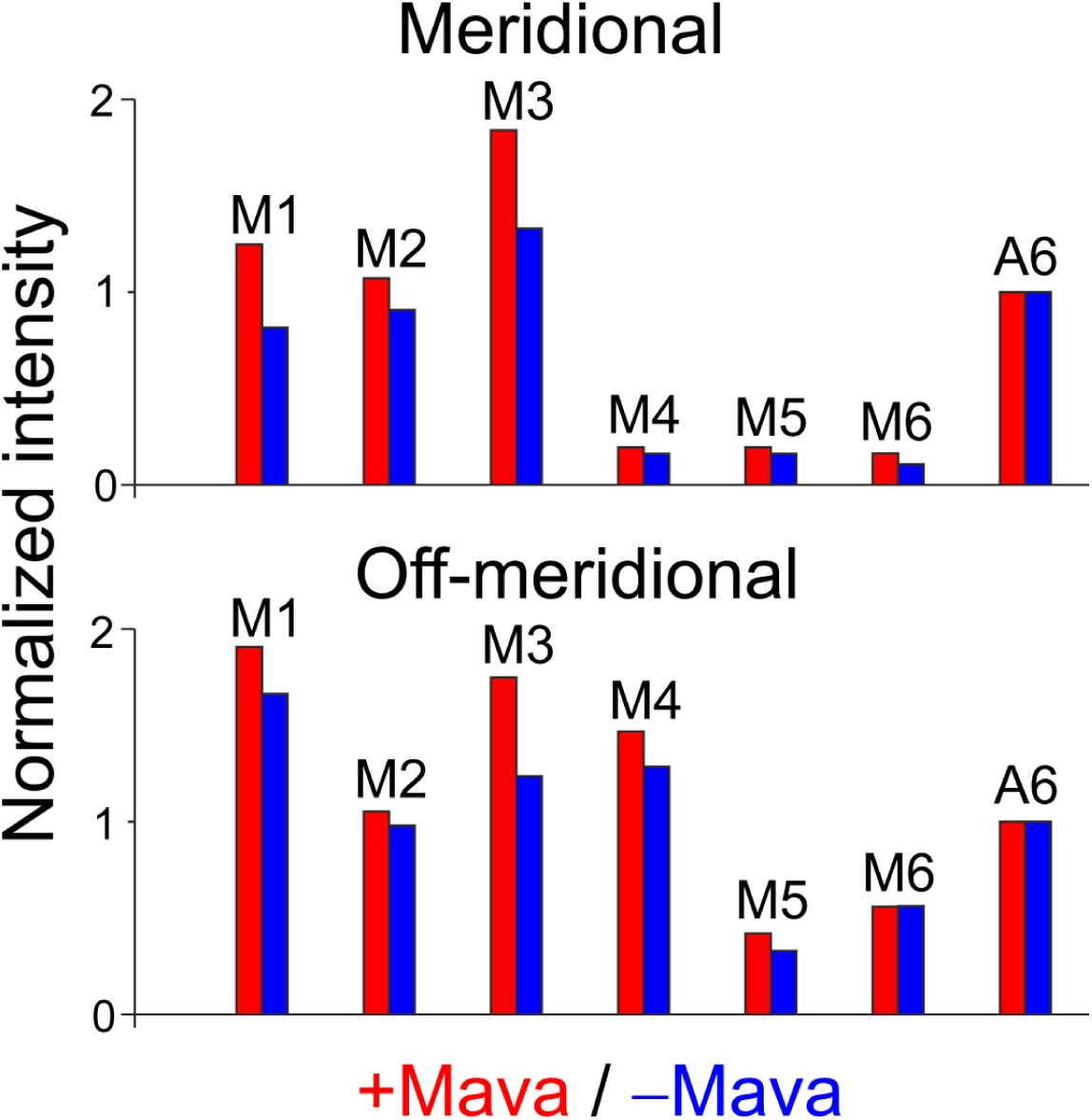

### Effect of mavacamten on the meridional (upper panel) and off-meridional (lower panel) intensities of myosin layer lines

Because the diffraction patterns were collected from different samples, the meridional (reciprocal radius range 0 – 0.0015 Å) and off-meridional (0.0015 – 0.015 Å) intensities were normalized to that of the actin A6 layer line, presumed to be unaffected by the myosin-specific drug mavacamten. The normalized intensities of myosin layer lines M1 – M6 in the absence of mavacamten (blue) were lower than those collected in its presence (red). The reduction in the normalized off-meridional intensities was up to 29%.

